# Understanding the factors that shape patterns of nucleotide diversity in the house mouse genome

**DOI:** 10.1101/275610

**Authors:** Tom R. Booker, Peter D. Keightley

## Abstract

A major goal of population genetics has been to determine the extent to which selection at linked sites influences patterns of neutral nucleotide diversity in the genome. Multiple lines of evidence suggest that diversity is influenced by both positive and negative selection. For example, in many species there are troughs in diversity surrounding functional genomic elements, consistent with the action of either background selection (BGS) or selective sweeps. In this study, we investigated the causes of the diversity troughs that are observed in the wild house mouse genome. Using the unfolded site frequency spectrum (uSFS), we estimated the strength and frequencies of deleterious and advantageous mutations occurring in different functional elements in the genome. We then used these estimates to parameterize forward-in-time simulations of chromosomes, using realistic distributions of functional elements and recombination rate variation in order to determine if selection at linked sites can explain the observed patterns of nucleotide diversity. The simulations suggest that BGS alone cannot explain the dips in diversity around either exons or conserved non-coding elements (CNEs). A combination of BGS and selective sweeps, however, can explain the troughs in diversity around CNEs. This is not the case for protein-coding exons, where observed dips in diversity cannot be explained by parameter estimates obtained from the uSFS. We discuss the extent to which our results provide evidence of sweeps playing a role in shaping patterns of nucleotide diversity and the limitations of using the uSFS for obtaining inferences of the frequency and effects of advantageous mutations.

**Author Summary:** We present a study examining the causes of variation in nucleotide diversity across the mouse genome. The status of mice as a model organism in the life sciences makes them an excellent model system for studying molecular evolution in mammals. In our study, we analyse how natural selection acting on new mutations can affect levels of nucleotide diversity through the processes of background selection and selective sweeps. To perform our analyses, we first estimated the rate and strengths of selected mutations from a sample of wild mice and then use our estimates in realistic population genetic simulations. Analysing simulations, we find that both harmful and beneficial mutations are required to explain patterns of nucleotide diversity in regions of the genome close to gene regulatory elements. For protein-coding genes, however, our approach is not able to fully explain observed patterns and we think that this is because there are strongly advantageous mutations that occur in protein-coding genes that we were not able to detect.

## Introduction

Starting with the discovery of a positive correlation between nucleotide polymorphism and the recombination rate in *Drosophila* in the late 1980s and early 1990s [1, 2], it has become clear that natural selection affects levels of genetic diversity across the genomes of many species [3, 4]. More recently, models incorporating selection at sites linked to those under observation have been shown to explain a large amount of the variation in diversity across the genome [5–8]. However, a persistent challenge has been to tease apart the contributions of positive and negative selection to the observed patterns.

Because the fates of linked alleles are non-independent, selection acting at one site may have consequences for variation and evolution at another. In broad terms, there are two models describing the effects of directional selection on neutral genetic diversity at linked sites, selective sweeps (SSWs) and background selection (BGS). SSWs occur when positively selected alleles spread through a population, dragging with them the haplotype on which they arose [9, 10]. There are a number of different types of SSW (reviewed in [11]), but in the present study, when not made explicit, we use the term selective sweep to refer to the effects of a single *de novo* advantageous mutation being driven to fixation by selection. BGS, on the other hand, occurs because the removal of deleterious mutations results in a loss of genetic diversity at linked neutral sites [12, 13]. The magnitudes of the effects of SSWs and BGS depend on the strength of selection, the rate of recombination and the mutation rate [10, 14, 15]. SSWs and BGS have qualitatively similar effects on genetic diversity, however, and many polymorphism summary statistics have little power to distinguish between them [12, 16].

Several studies have attempted to differentiate between BGS and SSWs. For example, Sattath *et al.* [17] examined patterns of nucleotide diversity around recent nucleotide substitutions in *Drosophila simulans*. Averaging across the entire genome, they observed a trough in diversity around nonsynonymous substitutions, whereas diversity was relatively constant around synonymous ones. This difference is expected under a model of recurrent SSWs, but not under BGS. Their results provide evidence that SSWs have been frequent in *D. simulans* since the species shared a common ancestor with *Drosophila melanogaster* (the outgroup used in that study). Similar results have been reported for *Capsella grandiflora* [18]. In humans [19], house mice [20] and maize [21], however, there is very little difference between the patterns of diversity around putatively neutral and potentially adaptive substitutions. These results have been interpreted as evidence that hard SSWs are infrequent in those species. However, Enard *et al.* [22] argued that since most adaptive substitutions are expected to occur in regions with the lowest functional constraint (and thus weaker BGS effects), the results of the Sattath test may be difficult to interpret in species with genomes that exhibit highly variable levels of functional constraint, such as humans and mice (but see [21]). Indeed, Enard *et al.* [22] found evidence that adaptive substitutions are fairly frequent in both protein-coding and non-coding portions of the human genome, suggesting that SSWs are common.

There are a number of methods that estimate the frequency and strength of advantageous mutations from models of the effects of selection at linked sites [11]. Recently, Elyashiv *et al.* [5] produced a map of the expected nucleotide diversity in *D. melanogaster* by fitting a model incorporating both BGS and hard SSWs to the genome-wide patterns of genetic diversity and the divergence between *D. melanogaster* and *D. simulans*. They concluded that sweeps are required to explain much of the genome-wide variation in diversity. However, the estimate of the deleterious per site mutation rate they obtained far exceeded published values of the point mutation rate in *D. melanogaster*. They, reasonably, attributed this discrepancy to the effects of selection at linked sites in addition to those they had explicitly modelled. The selection parameters estimated by Elyashiv *et al.* [5] were inferred from nucleotide diversity only. There is information in the distribution of allele frequencies, the site frequency spectrum (SFS), however, that can be used to estimate the distribution of fitness effects (DFE) for both deleterious and advantageous mutations [23–26]. In the present study, we estimate the DFE using such methods, and then use our estimates to parameterise the effects of BGS and SSWs.

In this study, we attempt to understand the influence of natural selection on variation at linked sites in the house mouse, *Mus musculus*. Specifically, we analyse *M. m. castaneus,* a sub-species which has been estimated to have a long-term effective population size (*N*_*e*_) of around 500,000 [27, 28], making it a powerful system in which to study molecular evolution in mammals. Both protein-coding genes and phylogenetically conserved non-coding elements (CNEs, which have roles in the regulation of gene expression [29]) exhibit signatures of natural selection in *M. m. castaneus* [20]. In particular, Halligan *et al.* [20] showed that there are substantial reductions in diversity surrounding protein-coding exons and CNEs, consistent with selection reducing diversity at linked sites. The trough in diversity surrounding exons was found to be ∼10x wider than the trough surrounding CNEs, suggesting that selection is typically stronger on protein sequences than regulatory sequences. However, Halligan *et al.* [20] found that troughs in diversity around recent nonsynonymous and synonymous substitutions in *M. m. castaneus* were similar. Taken at face value, this could be taken as evidence that SSWs are infrequent, but, in addition, Halligan *et al.* [20] found that there are also troughs in diversity around randomly chosen synonymous or nonsynonymous sites that are similar to those observed around substitutions. These results, therefore, suggest that selection at linked sites affects nucleotide diversity across large portions of the genome, making the analysis of patterns of diversity around substitutions difficult to interpret. Our understanding of the forces that have shaped patterns of diversity in the house mouse and mammals in general is, thus, somewhat unclear.

We analyse data on wild-caught *M. m. castaneus* individuals to obtain estimates of the distribution of fitness effects (DFEs) for several classes of functional elements in the mouse genome and then use these to parameterise forward-in-time simulations. We analyse several aspects of our simulation data: 1) the patterns of genetic diversity and the distribution of allele frequencies around both protein-coding exons and conserved non-coding elements; 2) the rates of substitution in different functional elements; and 3) the patterns of diversity around nonsynonymous and synonymous substitutions.

## Materials and Methods

### Samples and polymorphism data

We analysed the genome sequences of 10 wild-caught *M. m. castaneus* individuals sequenced by Halligan *et al.* [20]. The individuals were sampled from an area that is thought to include the ancestral range of the species [28]. A population structure analysis suggested that the individuals chosen for sequencing came from a single randomly mating population [27]. Sampled individuals were sequenced to an average depth of ∼30x using Illumina technology. Reads were mapped to version mm9 of the mouse genome and variants called as described in Halligan *et al.* [20]. Only single nucleotide polymorphisms were considered, and insertion/deletion polymorphisms were excluded from downstream analyses. We used the genome sequences of *Mus famulus* and *Rattus norvegicus* as outgroups in this study. For *M. famulus*, a single individual was sequenced to high coverage and mapped to the mm9 genome [20]. For *R. norvegicus*, we used the whole genome alignment of the mouse (mm9) and rat (rn4) reference genomes from UCSC.

For the DFE-alpha analysis (*see below*), the underlying model assumes a single, constant mutation rate. Hypermutable CpG sites strongly violate this assumption, so CpG-prone sites were excluded as a conservative way to remove CpG sites from our analyses. A site was labelled as CpG-prone if it is preceded by a C or followed by a G in the 5’ to 3’ direction in either *M. m. castaneus, M. famulus* or *R. norvegicus*. Additionally, sites that failed a Hardy-Weinberg equilibrium test (*p* < 0.002) were excluded from further analysis, because they may represent sequencing errors.

### Functional elements in the murid genome

In this study, we considered three different classes of functional elements in the genome: the exons and untranslated regions (UTRs) of protein-coding genes and conserved non-coding elements (CNEs).

Coordinates for canonical splice-forms of protein-coding gene orthologs between *Mus musculus* and *Rattus norvegicus* were obtained from version 67 of the Ensembl database. We used these to identify untranslated regions (UTRs) as well as 4-fold and 0-fold degenerate sites in the coding regions. We made no distinction between 3’ and 5’ UTRs in the analysis. Genes containing alignment gaps affecting >80% of sites in either outgroup and genes containing overlapping reading frames were excluded. This left a total of 18,171 autosomal protein-coding genes.

The locations of conserved non-coding elements (CNEs) in the house mouse genome were identified as described by Halligan *et al.* [20].

Estimating the parameters of the distribution of fitness effects (DFE) for a particular class of sites using DFE-alpha (*see below*) requires neutrally evolving sequences for comparison. When analysing 0-fold degenerate sites and UTRs, we used 4-fold degenerate sites as the comparator. For CNEs, we used non-conserved sequence in the flanks of CNEs. Halligan *et al.* [20] found that, compared to the genome-wide average, nucleotide divergence between mouse and rat in the ∼500bp on either side of CNEs is ∼20% lower than that of intergenic DNA distant from CNEs, suggesting functional constraint in these regions. For the purpose of obtaining a quasi-neutrally evolving reference class of sequence and to avoid these potentially functional sequences, we therefore used sequence flanking the edges of each CNE, offset by 500bps. For each CNE, the total amount of flanking sequence used in the analysis was equal to the length of the focal CNE, split evenly between the upstream and downstream regions. CNE-flanking sequences overlapping with another annotated feature (i.e. exon, UTR or CNE) or the flanking sequence of another CNE were excluded.

### The site frequency spectrum around functional elements

For distances of up to 100Kbp on either side of exons and 5Kbp on either side of CNEs, the non-CpG-prone sites in non-overlapping windows of 1Kbp and 100bp, respectively, were extracted. Sites within analysis windows that overlapped with any of the annotated features described above, or that contained missing data in *M. m. castaneus* or either outgroup were excluded. The data for analysis windows were collated based on the distance to the nearest CNE or exon, from which we calculated nucleotide diversity and Tajima’s *D*.

### Overview of DFE-alpha analysis

The distribution of allele frequencies in a sample, referred to as the site frequency spectrum (SFS), provides information on evolutionary processes. Under neutrality the SFS reflects past demographic processes, such as population expansions and bottlenecks, and potentially the effects of selection at linked sites. The allele frequency distribution will also be distorted if focal sites are subject to functional constraints. The SFS therefore contains information on the strengths and frequencies of mutations with different selective effects, known as the distribution of fitness effects (hereafter the DFE). Note that balancing selection may maintain alleles at intermediate frequencies [30], but we assume that the contribution of this form of selection to overall genomic diversity is negligible.

DFE-alpha estimates selection parameters using information contained in the SFS by a two-step procedure [24]. First, a demographic model is fitted to data for a class of putatively neutral sites. Conditional on the demographic parameter estimates, the DFE is then estimated for the selected sites. In the absence of knowledge of ancestral or derived alleles, the ‘folded’ SFS can be used to estimate the demographic model and the DFE for harmful variants (hereafter referred to as the dDFE) [24]. If information from one or more outgroup species is available, and the ancestral state for a segregating site can be inferred, one can construct the ‘unfolded’ SFS (uSFS). In the presence of positive selection, such that advantageous alleles segregate at an appreciable frequency, the parameters of the distribution of fitness effects for advantageous mutations can be estimated from the uSFS [25, 26, 31]. In this study, we estimate the proportion of new mutations occurring at a site that are advantageous (*p*_*a*_) and the strength of selection acting on them (*N*_*e*_*s*_*a*_).

### Inference of the uSFS and the DFE

We inferred the distributions of derived allele frequencies in our sample for 0-fold and 4-fold sites, UTRs, CNEs and CNE-flanks using *M. famulus* and *R. norvegicus* as outgroups, using the two-outgroup method implemented in ml-est-sfs v1.1 [31]. This method employs a two-step procedure conceived to address the biases inherent in parsimony methods. The first step estimates the rate parameters for the tree under the Jukes-Cantor model by maximum likelihood assuming a single mutation rate. Conditional on the rate parameters, the individual elements of the uSFS are then estimated.

DFE-alpha fits discrete population size models, allowing up to two changes in population size through time. For each class of putatively neutral sites, one-, two-and three-epoch models were fitted by maximum likelihood and the models with the best fit (as judged by likelihood ratio tests) were used in further analyses. When fitting the three-epoch model, we ran DFE-alpha (v2.16) 10 times with a range of different search algorithm starting values, in order to check convergence.

In the cases of 4-fold sites and CNE-flanks, the inferred uSFSs exhibited a higher proportion of high frequency derived alleles than expected under the best-fitting demographic model (Figure S1) (hereafter referred to as an uptick). Such an increase is not possible under the single population, single locus demographic models assumed. There are several possible explanations for the uptick: 1) mis-inference of the uSFS due to an inadequacy of the model assumed in ml-est-sfs; 2) failure to capture the demographic history of *M. m. castaneus* by the models implemented in DFE-alpha; 3) sequencing errors in *M. m. castaneus* or either outgroup generating spurious signals of divergence; 4) SSWs, since they can drag linked alleles to high frequencies [32, 33]; 5) cryptic population sub-division in our sample of mouse individuals; and positive selection, acting on the putatively neutral sites themselves. We think this latter explanation is unlikely, however, since there is little evidence for selection on synonymous codon usage in *Mus musculus* [34]. With the exception of direct selection affecting the putatively neutral class of sites, the above sources of bias should also affect the selected class of sites [31, 35, 36]. We therefore corrected the selected sites uSFS prior to inferring selection parameters by subtracting the proportional deviation between the neutral uSFS expected under the best-fitting demographic model and the observed neutral uSFS (following Keightley *et al.* [31]; see Supplementary Methods).

Simultaneous inference of the DFE for harmful mutations (dDFE) and adaptive mutation parameters was performed using DFE-alpha (v.2.16) [25]. A gamma distribution has previously been used to model the dDFE, since it can take a variety of shapes and has only two parameters [37]. However, more parameter-rich discrete point mass distributions provide a better fit to nonsynonymous polymorphism site data in wild house mice [38]. We therefore compared the fit of one, two and three discrete class dDFEs and the gamma distribution, and also included one or more classes of advantageous mutations. Nested DFE models were compared using likelihood ratio tests, and non-nested models were compared using Akaike’s Information Criteria (AIC). Goodness of fit was also assessed by comparing observed and expected uSFSs using the *χ*^2^-statistic, but the numbers of sites in the *i*^*th*^ and *n-i*^*th*^ classes are non-independent, so formal hypothesis tests were not performed.

We constructed profile likelihoods to obtain confidence intervals. Two unit reductions in *logL*, on either side of the maximum likelihood estimates (MLEs) were taken as approximate 95% confidence limits.

### Two methods for inferring the rates and effects of advantageous mutations based on the uSFS

It has been suggested that estimates of the DFE obtained based on the uSFS may be biased if sites fixed for the derived allele are included in calculations [26]. Sites fixed for the derived allele are typically a frequent class in the uSFS, and therefore strongly influence parameter estimates. Bias can arise, for example, if the selection strength has changed since the split with the outgroup, such that the number of sites fixed for the derived allele do not reflect the selection regime that generated current levels of polymorphism. If nucleotide divergence and polymorphism are decoupled in this way, selection parameter estimated from only polymorphism data (and sites fixed for ancestral alleles) may therefore be less biased than those obtained when using the full uSFS. To investigate this possibility, we estimated selection parameters either utilising the full uSFS (we refer to this method as Model A) or by analysing the uSFS while fitting an additional parameter (Supplementary Methods), such that sites fixed for the derived allele do not contribute to estimates of the selection parameters (we refer to this method as Model B).

Certain alleles present in a sample of individuals drawn from a population may appear to be fixed that are, in fact, polymorphic. Attributing such polymorphisms to between-species divergence may then influence estimates of the DFE by increasing the number of sites fixed for the derived allele (note that this would only affect estimates obtained under Model A). We corrected the effect of polymorphism attributed to divergence using an iterative approach as follows. When fitting selection or demographic models, DFE-alpha produces a vector of expected allele frequencies. Using this vector, we inferred the expected proportion of polymorphic sites that appear to be fixed for the derived allele. This proportion was then subtracted from the fixed derived class and distributed among the polymorphism bins according to the allele frequency vector. We then refitted the model using this corrected uSFS, and this procedure was applied iteratively until convergence (See Supplementary Methods). For each site class, convergence was achieved within five iterations and the selection parameters for each class did not substantially change between iterations.

### Forward-in-time simulations modelling background selection and selective sweeps

We performed forward-in-time simulations in SLiM v1.8 [39] to assess whether the observed patterns of diversity around functional elements [20] can be explained by SSWs or BGS caused by mutations originating in the elements themselves. These simulations focussed on either protein-coding exons or CNEs. We also ran SLiM simulations to model the accumulation of between-species divergence under our estimates of the DFE. In all our simulations, we either assumed the estimates of selection parameters obtained from the full uSFS (Model A) or those obtained when sites fixed for the derived allele do not contribute to parameter estimates (Model B).

Models of BGS and recurrent SSWs predict that the magnitudes of their effects are sensitive to the rate of recombination and mutation rate and the strength of selection [14, 40, 41]. To parameterise our simulations, we used estimates of compound parameters scaled by *N*_*e*_. For example, estimates of selection parameters obtained from DFE-alpha are expressed in terms of *N*_*e*_*s* (where *s* is the difference in fitness between homozygotes for ancestral and derived alleles, assuming semi-dominance). For a population where *N*_*e*_ = 1,000 and *s* = 0.05, for example, the strength of selection is therefore approximately equivalent to that of a population where *N*_*e*_ = 10,000 and *s* = 0.005. By scaling parameter values according to the population size of the simulations (*N*_*sim*_), we modelled the much larger *M. m. castaneus* population (*N*_*e*_ ≅ 500,000 [42] in a computationally tractable way.

#### 1. Annotating simulated chromosomes

Functional elements are non-randomly distributed across the house mouse genome. For example, protein-coding exons are clustered into genes and CNEs are often found close to other CNEs [20]. Incorporating this distribution into simulations is important when modelling BGS and recurrent SSWs, because their effects on neutral diversity depend on the density of functional sequence [14, 43]. We incorporated the distribution as follows. For each simulation replicate, we chose a random position on an autosome, which was itself randomly selected (with respect to length). The coordinates of the functional elements (exons, UTRs and CNEs) in the 500Kbp downstream of that position were used to annotate a simulated chromosome of the same length. For simulations focussing on exons (CNEs), we only used chromosomal regions that had at least one exon (CNE).

#### 2. Mutation, recombination and selection in simulations

We used an estimate of the population scaled mutation rate, *θ=4N*_*e*_*μ*, to set the mutation rate (*μ*) in simulations, such that levels of neutral polymorphism approximately matched those of *M. m. castaneus*. Diversity at putatively neutral sites located close to functional elements (for example, 4-fold synonymous sites) may be affected by BGS and SSWs. To correct for this, we used an estimate of *θ* = 0.0083, based on the average nucleotide diversity at non-CpG-prone sites at distances >75Kbp from protein-coding exons. This distance was used, because it the approximate distance beyond which nucleotide diversity remains flat. The mutation rate in simulations was thus set to 0.0083/*4N*_*sim*_.

Variations in the effectiveness of selection at linked sites, due to variation in the rate of recombination across the genome, may not be captured by simulations that assume a single rate of crossing over. Recently, we generated a map of variation in the rate of crossing-over for *M. m. castaneus* using a coalescent approach [44], quantified in terms of the population scaled recombination rate *ρ=4N*_*e*_*r*. Recombination rate variation in the 500Kbp region used to obtain functional annotation was used to specify the genetic map for individual simulations.

We modelled natural selection at sites within protein-coding exons, UTRs and CNEs in the simulations using the estimates of selection parameters obtained from the DFE-alpha analysis. In the case of protein-coding exons, 25% of sites were set to evolve neutrally (i.e. synonymous sites), and the fitness effects of the remaining 75% were drawn from the DFE inferred for 0-fold sites (hereafter termed nonsynonymous sites in the simulations). For mutations in UTRs and CNEs, 100% were drawn from the DFEs inferred for those elements. Population scaled selection coefficients were divided by *N*_*sim*_ to obtain values of *s* for use in simulations. All selected mutations were assigned a dominance coefficient of 0.5, as assumed by DFE-alpha.

#### 3. Patterns of diversity around functional elements in simulations

We examined the contributions of BGS and recurrent SSWs to the troughs in diversity observed around protein-coding exons and CNEs using forward-in-time simulations. Focussing on either protein-coding exons or CNEs, we performed three sets of simulations. The first incorporated only harmful mutations (causing BGS), the second only advantageous mutations (causing SSWs), and the third set incorporated both (causing both processes). Thus, under a given set of DFE estimates, we performed six sets of simulations (three sets focussing on exons and three sets focussing on CNEs). For each simulation set, 2,000 SLiM runs were performed, each using a randomly sampled 500Kbp region of the genome. In each SLiM run, populations of *N*_*sim*_= 1,000 diploid individuals were allowed to evolve for 10,000 generations (10*N*_*sim*_) in order to approach mutation-selection-drift balance. At this point, 200 randomly chosen haploid chromosomes were sampled from the population and used to construct SFSs.

For each set of simulations, segregating sites in windows surrounding functional elements were analysed in the same way as for the *M. m. castaneus* data (*see above*). The SFSs for all windows at the same distance from an element were collated. Analysis windows around protein-coding exons were oriented with respect to the strand orientation of the actual gene. Neutral sites near the tips of simulated chromosomes only experience selection at linked sites from one direction, so analysis windows located within 60Kbp of either end of a simulated chromosome were discarded. For a given distance to a functional element, we obtained confidence intervals around individual statistics by bootstrapping analysis window 1,000 times.

Mutation rate variation is expected to contribute to variation in nucleotide diversity. Nucleotide divergence between mouse and rat is relatively constant in the intergenic regions surrounding protein-coding exons [20], suggesting that mutation rate variation is not responsible for the troughs in diversity around exons. Around CNEs, however, there is a pronounced dip in nucleotide divergence between *M. m. castaneus* and the rat. A likely explanation for this is that alignment-based approaches to identify CNEs fail to identify the edges of some elements, resulting in the inclusion of functionally constrained sequence in the analysis windows close to CNEs. This factor was not incorporated in our simulations, so in order to correct for this constraint, allowing us to compare diversity around CNEs in *M. m. castaneus* with our simulation data, we scaled values as follows. We divided nucleotide diversity by between-species divergence, in this case mouse-rat divergence, giving a statistic (*π/d*_*rat*_) that reflects diversity corrected for mutation rate variation. We then multiplied the *π/d*_*rat*_ values by the mean mouse-rat divergence in regions further than 3Kbp from the edges of CNEs to obtain values on the same scale as our simulation data.

When comparing the patterns of diversity around functional elements in our simulations with the observations from *M. m. castaneus*, we used the root mean square (RMS) as a measure of goodness-of-fit.

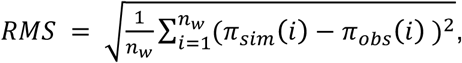

where *π*_*sim*_*(i)* and *π*_*obs*_*(i)* are the diversity values from simulations and *M. m. castaneus*, respectively, in window *i* around a particular class of functional element and *n*_*w*_ is the total number of analysis windows. Approximate confidence intervals for RMS values were obtained using the bootstrap replicates described above.

#### 4. Re-inferring the DFE based on simulated population data

We performed two additional sets of simulations to model the accumulation of between-species nucleotide divergence under the DFE estimates obtained by analysis of the full uSFS (i.e. Model A) and those obtained when sites fixed for the derived allele did not contribute to selection parameters (i.e. Model B). These simulations were the same as those described above, except that we ran them for additional generations to approximate the mouse-rat divergence. We ran 4,000 replicates of these simulations. Using polymorphic sites and sites fixed for the derived allele, we constructed the uSFS for each class of functional sites.

In order to model the mouse-rat divergence, we required a time frame to approximate the neutral divergence between those two species. Neutral divergence between *M. m. castaneus* and *R. norvegicus* (*K*_*rat*_) is ∼15% at non-CpG-prone sites far from protein-coding exons. Under neutrality, divergence is expected to be equal to *2Tμ*, where *T* is the time in generations since the two-species shared a common ancestor and *μ* is the mutation rate per base pair per generation. In the simulations, the mutation rate was 2.075 × 10^−6^ bp^−1^ (recall that we scaled mutations rates using an estimate of *4N*_*e*_*µ*) and since *K*_*rat*_ = 0.15, *T* = 36,145 generations. We thus ran simulations incorporating both deleterious and advantageous mutations, focussing on exons, for 46,145 generations, discarding the first 10,000 as burn-in. At the final generation, we constructed the uSFS for synonymous and nonsynonymous sites from 20 randomly sampled haploid chromosomes. To obtain a proxy for mouse-rat divergence, we counted all substitutions that occurred after the 10*N*_*sim*_ burn-in phase plus any derived alleles present in all 20 haploid chromosomes.

Using the uSFSs for synonymous and nonsynonymous sites obtained from the simulations, we estimated selection parameters using the methods described above. We first fitted one-, two-and three-epoch demographic models to simulated synonymous site data. For the simulations assuming Model A or Model B, we found that the three-epoch demographic model gave the best fit to the simulated synonymous site uSFS in both cases. Using the expected uSFS under the three-epoch model, we performed the demographic correction (Supplementary Methods) before estimating selection parameters. When estimating selection parameters based on simulation data, we used the same methods as used for the analysis of the *M. m. castaneus* data, i.e. the DFE for Model A simulations was estimated using Model A *etc.*

#### 5. Patterns of diversity around recent nonsynonymous and synonymous substitutions

Comparisons of the average level of nucleotide diversity around recent synonymous and nonsynonymous substitutions have been used to test for positive selection [17–21]. In *M. m. castaneus* there is essentially no difference in diversity around recent substitutions at 0-fold and 4-fold sites [20]. This could reflect a paucity of SSWs, or alternatively, this particular test may be unable to discriminate between BGS and SSWs in mice. Using our simulation data, in which SSWs are relatively frequent, we tested whether patterns of diversity around selected and neutral substitutions reveals the action of positive selection. In their study, Halligan *et al.* [20] used *M. famulus* as an outgroup to locate recent substitutions, because it is much more closely related to *M. musculus* than the rat. We obtained the locations of nucleotide substitutions in our simulations as follows. Neutral divergence between *M. m. castaneus* and *M. famulus* (*K*_*fam*_) is 3.4%. In the simulations, given that the mutation rate was 2.075×10^−6^, 8,193 generations are sufficient to approximate the *M. m. castaneus* lineage since its split with *M. famulus K*_*fam*_. Thus, all substitutions that occurred in 8,193 generations were analysed. Neutral diversity around synonymous and nonsynonymous substitutions in non-overlapping windows of 1,000bp up to 100Kbp from substituted sites were then extracted from the simulations. Sites in analysis windows that overlapped with functional elements were excluded. If two substitutions of the same type were located less than 100Kbp apart, analysis windows extended only to the midpoint of the two sites.

Except where noted, all analyses were conducted using custom Python and Perl scripts (available on request).

## Results

To investigate genetic variation around functional elements in house mice, we analysed the genomes of 10 wild-caught individuals that had been sequenced to high coverage [20]. We compared nucleotide polymorphism and between-species divergence in three classes of functional sites (0-fold sites, UTRs and CNEs) with polymorphism and divergence at linked, putatively neutral sequences (4-fold sites and CNE-flanks). The three classes of functional sites had lower levels of within-species polymorphism and between-species divergence than their neutral comparators (Table 1). This is the expected pattern if natural selection keeps deleterious alleles at low frequencies, preventing them from reaching fixation. Tajima’s *D* is more negative for 0-fold sites, UTRs and CNEs than for their neutral comparators (Table 1), further indicating the action of purifying selection in those classes of sites. It is notable that the two neutral site types exhibited negative Tajima’s *D*, indicating that rare variants are more frequent than expected in a Wright-Fisher population (Table 1). This is consistent either with a recent population expansion or the widespread effects of selection on linked sites, both of which may be relevant for this population [20, 44].

**Table 1.** Summary statistics for five classes of sites in *M. m. castaneus.* All values refer to non-CpG prone sites. Nucleotide divergences between *M. m. castaneus* and *M. famulus* (*d*_*fam*_) and between *M. m. castaneus* and *R. norvegicus* (*d*_*rat*_) were estimated by maximum likelihood using the method described in [31].

| | $\pi$ (%) | Tajima's $D$ | $d_{fam}$ (%)* | $d_{rat}$ (%)* | Sites (Mb) |
| --- | --- | --- | --- | --- | --- |
| *0-fold* | 0.134 | -0.763 | 0.239 | 2.93 | 10.2 |
| *4-fold* | 0.628 | -0.627 | 1.06 | 12.7 | 1.49 |
| *CNE* | 0.274 | -1.03 | 0.418 | 3.67 | 24.6 |
| *CNE flank* | 0.670 | -0.602 | 1.03 | 13.8 | 17.8 |
| *UTR* | 0.438 | -0.702 | 0.802 | 10.0 | 11.3 |

### Inferring the unfolded site frequency spectrum

The distribution of derived allele frequencies in a class of sites (the unfolded site frequency spectrum-uSFS) potentially contains information on the frequency and strength of selected mutations. We estimated the uSFSs for 0-fold sites, UTRs and CNEs using a probabilistic method incorporating information from two outgroup species [31]. This method attempts to correct for biases that are inherent in parsimony methods.

A population’s demographic history is expected to affect the shape of the SFS. DFE-alpha attempts to correct this by fitting a population size change model to the neutral site class, and, conditional on the estimated demographic parameters, estimates the DFE for linked, selected sites. In the case of 4-fold sites and CNE flanks, a 3-epoch model provided the best fit to the data, based on likelihood ratio tests (Table S1) The trajectories of the inferred population size changes were similar in each case, i.e. a population bottleneck followed by an expansion (Table S2). However, the magnitude of the changes and the duration of each epoch differed somewhat (Table S2). A possible explanation is that the demographic parameter estimates are affected by selection at linked sites, which differs between site classes [45–47].

We found that the 4-fold site and CNE-flank uSFSs exhibited an excess of high frequency derived alleles relative to expectations under the best-fitting neutral demographic models (Figure S1). For example, *χ*^2^-statistics for the difference between the observed and fitted number of sites for the last uSFS element (i.e. 19 derived alleles) were 245.9 and 505.6 for 4-fold sites and CNE-flanks, respectively. It is reasonable to assume that the differences between fitted and observed values are caused by processes that similarly affect the linked selected site class. We therefore corrected the 0-fold, UTR and CNE uSFSs by subtracting the proportional deviations between fitted and observed values for neutral site uSFSs prior to estimating selection parameters (see Supplementary Methods). Applying this correction (hereafter referred to as the demographic correction) appreciably reduced the proportion of high frequency derived variants (Figure 1).

**Figure 1.**
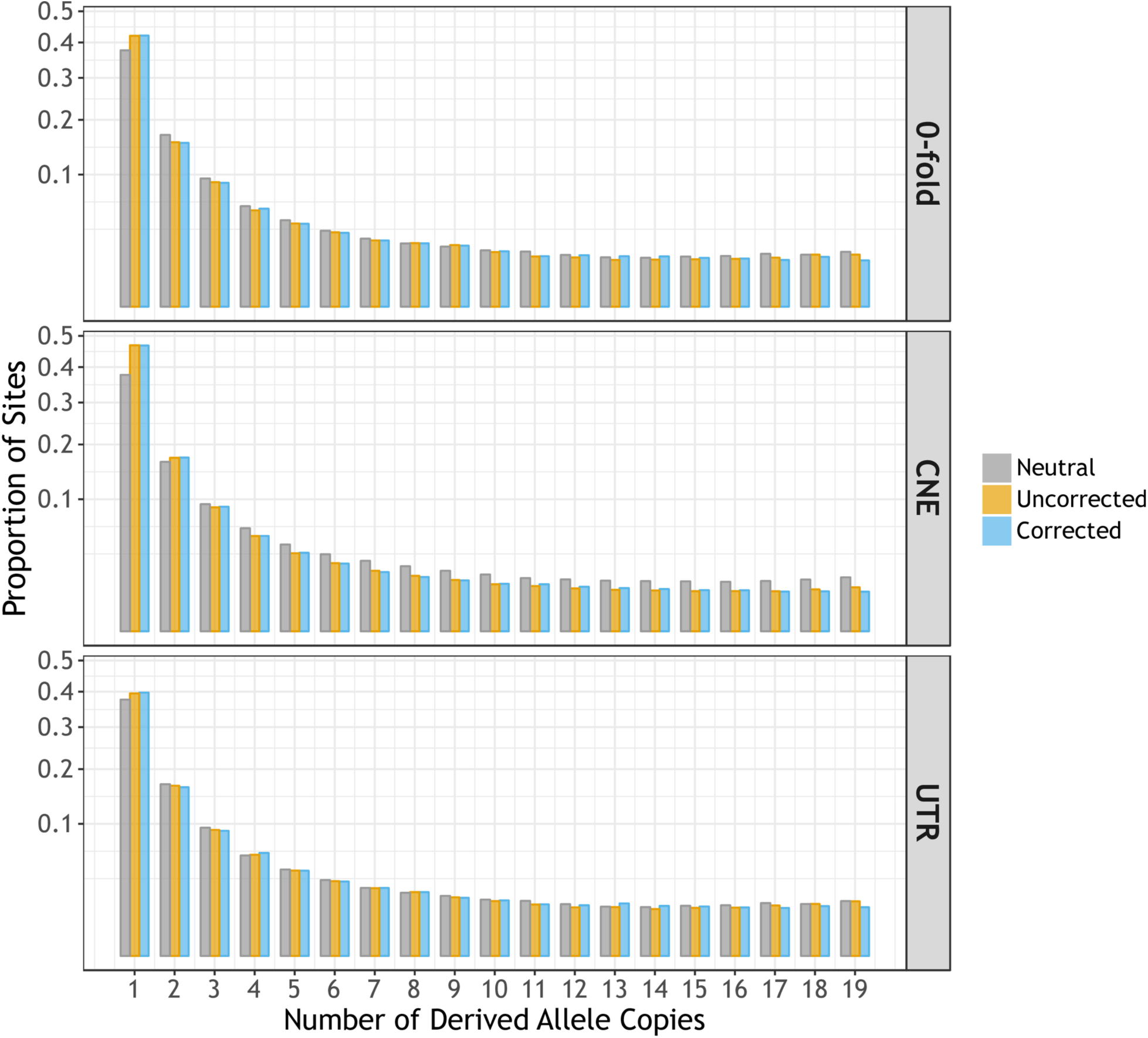
The uSFS for three classes of functional sites (yellow and blue bars) compared to a putatively neutral comparator (grey bars). The neutral comparator for 0-fold sites and UTRs was 4-fold degenerate synonymous sites in both cases. For CNEs, the neutral comparator was CNE-flanking sequence. The expected uSFS under a demographic model fitted to a neutral comparator was used to correct the uSFS for the corresponding selected sites (see *Methods*).

### Estimating the frequencies and strengths of deleterious and advantageous mutations

We inferred the DFE for harmful mutations (dDFE) and the rate and strength of advantageous mutations based on the uSFSs for the three different classes of functional sites using DFE-alpha under two different models (Table 2). The first, as described by Schneider *et al.* [25], makes use of the full uSFS, including sites fixed for the derived allele (hereafter Model A). The second (hereafter Model B), incorporated an additional parameter that absorbs the contribution of sites fixed for the derived allele (see Supplementary Methods). This was motivated by the possibility that between-species divergence may be decoupled from within-species polymorphism (e.g. due to changing selection regimes), and this could lead to spurious estimates of selection parameters [26, 48]. Since Model A is nested within Model B, the two can be compared using likelihood ratio tests. In the remainder of the study, results obtained under Model A are shown in parallel with results obtained under Model B.

**Table 2.** Parameter estimates for the distribution of fitness effects for three classes of sites in *M. m. castaneus* obtained under two models. The first (Model A) estimates of selection parameters based on the full uSFS. Under the second method (Model B), sites fixed for the derived allele were prevented from influencing estimates of selection parameters. The bracketed values are 95% confidence intervals obtained from profile likelihoods. The parameters shown are: *p*_*i*_ = the proportion of mutations falling into the *i*^*th*^ deleterious class; *N*_*e*_*s*_*i*_ = the scaled homozygous selection coefficient of the *i*^*th*^ deleterious class; *p*_*a*_ = the proportion of advantageous mutations; *N*_*e*_*s*_*a*_ = the scaled homozygous selection coefficient of the advantageous mutation class.

| **Model A: DFE inferred from the full uSFS** |  |  |  |
| --- | --- | --- | --- |
|  | **0-fold** | **UTR** | **CNE** |
| $N_e s_1$ | -0.045 | -0.097 | -0.323 |
| $p_1$ | 0.191 | 0.701 | 0.352 |
| $N_e s_2$ | -104 | -39.1 | -3.98 |
| $p_2$ | 0.806 | 0.286 | 0.278 |
| $N_e s_3$ | - | - | -77.9 |
| $p_3$ | - | - | 0.360 |
| $N_e s_a$ | 7.27<br>[4.62 – 11.7] | 5.32<br>[3.91 – 7.03] | 9.17<br>[7.00 – 20.9] |
| $p_a$ | 0.0030<br>[0.0019 – 0.0048] | 0.013<br>[0.0097 – 0.019] | 0.0098<br>[0.0037 – 0.0099] |
| **Model B: Sites fixed for the derived allele do not contribute to parameter estimates** |  |  |  |
|  | **0-fold** | **UTR** | **CNE** |
| $N_e s_1$ | -0.171 | -0.160 | -0.253 |
| $p_1$ | 0.184 | 0.689 | 0.342 |
| $N_e s_2$ | -100 | -32.0 | -3.84 |
| $p_2$ | 0.806 | 0.281 | 0.286 |
| $N_e s_3$ | - | - | -76.3 |
| $p_3$ | - | - | 0.365 |
| $N_e s_a$ | 8.30<br>[6.24 – 10.1] | 6.96<br>[5.53 – 8.69] | 8.60<br>[4.37 – 12.6] |
| $p_a$ | 0.010<br>[0.0030 – 0.0183] | 0.0294<br>[0.0181 – 0.0436] | 0.008<br>[0.0004 – 0.0100] |

We performed a comparison of different DFE models, including discrete distributions that have one, two or three mutational effect classes and the gamma distribution including or not including advantageous mutations. For each class of functional sites, DFE models with several classes of deleterious mutational effects and a single class of advantageous effects gave the best fit (Table S3). For each class of functional sites, only a single class of advantageous mutations was supported, since additional classes of advantageous mutations did not significantly increase likelihoods (Table S4). This presumably reflects a lack of power. These best-fitting models were identified whether we estimated the DFE under Model A or Model B. Parameter estimates pertaining to the dDFE were also similar between Models A and B (Table 2).

In our current study, we estimated selection parameters based on the uSFS, whereas earlier studies on mice used the distribution of minor allele frequencies, i.e. the ‘folded’ SFS [20, 27, 49–51]. A possible consequence of using the folded SFS is that advantageous mutations segregating at intermediate to high frequencies are allocated to the mildly deleterious class. In the case of 0-fold sites, for example, the best-fitting DFE did not include mutations with scaled effects in the range of 1 < |*N*_*e*_*s*| < 100 (Table 2). This contrasts with previous studies using the folded SFS which found an appreciable proportion of mutations in the 1 < |*N*_*e*_*s*| < 100 range [20, 38]. Because this difference may have an effect on the reductions in diversity caused by background selection, we performed simulations incorporating either the gamma dDFEs inferred from analysis of the folded SFS by Halligan *et al.* [20] or the discrete dDFEs inferred in the present study (results below).

For all classes of functional sites, we inferred that moderately positively selected mutations are fairly frequent under both Models A and B (Table 2). In the case of 0-fold sites, for example, the frequency of advantageous mutations was 0.3% (under Model A). Across the three classes of sites, the average scaled selection strengths of advantageous mutations were fairly similar (Table 2), i.e. *N*_*e*_*s ∼* 8, implying that *s* is on the order of 10^−5^ (assuming *N*_*e*_ = 500,000; [42]). We found that estimates of the frequency of advantageous mutations (*p*_*a*_) obtained under Model B for 0-fold sites and UTRs were ∼3 times higher than those obtained under Model A. Confidence intervals overlapped, however (Table 2). In the cases of both 0-fold sites and UTRs, Model B fitted significantly better than Model A, as judged by likelihood ratio tests (0-fold sites, *χ*^2^_1 d.f._= 4.2; *p* = 0.04; UTRs, *χ*^2^_1 d.f._= 9.9; *p* = 0.002). Interestingly, in the case of CNEs, Models A and B did not differ significantly in fit (*χ*^2^_1 d.f._= 0.26; *p =* 0.60) and estimates of the advantageous mutation parameters were very similar (Table 2).

### Forward-in-time population genetic simulations

We conducted forward-in-time simulations to examine whether estimates of the DFE obtained by analysis of the uSFS predict patterns of diversity observed around functional elements. In our simulations, we used estimates of selection parameters obtained by DFE-alpha for 0-fold sites, UTRs and CNEs, assuming either Model A (i.e. from the full uSFS) or Model B (i.e. by absorbing the contribution of sites fixed for the derived allele with an additional parameter). The selection parameter estimates obtained under Models A and B resulted in major differences in the patterns of diversity around functional elements.

#### i) Patterns of nucleotide diversity around functional elements in simulated populations

Using the selection parameter estimates obtained from DFE-alpha (Table 2), we performed simulations incorporating deleterious mutations, advantageous mutations or both advantageous and deleterious. Our analysis involved computing diversity in windows surrounding functional elements and comparing the diversity patterns with those seen in *M. m. castaneus.* In order to aid visual comparisons, we divided nucleotide diversity (*π*) at all positions by the mean *π* at distances greater than 75Kbp and 4Kbp away from exons and CNEs, respectively. These distances were chosen as they are the approximate values beyond which *π* remains constant.

Simulations incorporating only deleterious mutations predicted a chromosome-wide reduction in genetic diversity. Around exons and CNEs diversity plateaued at values that were ∼94% of the neutral expectation (Figures S2–3). However, simulations involving only BGS did not fully predict the observed troughs in diversity around functional elements. The predicted troughs in diversity around both protein-coding exons and CNEs, were not as wide nor as deep as those observed in the real data (Figures 2–3). Similar predictions were obtained for Models A or B (Figures 2–3) or for the gamma dDFEs inferred by Halligan *et al.* [20] (Figure S4). Our simulations incorporating deleterious mutations suggest, then, that while BGS affects overall genetic diversity across large portions of the genome, but positive selection presumably also makes a substantial contribution to the dips in diversity around functional elements.

**Figure 2.**
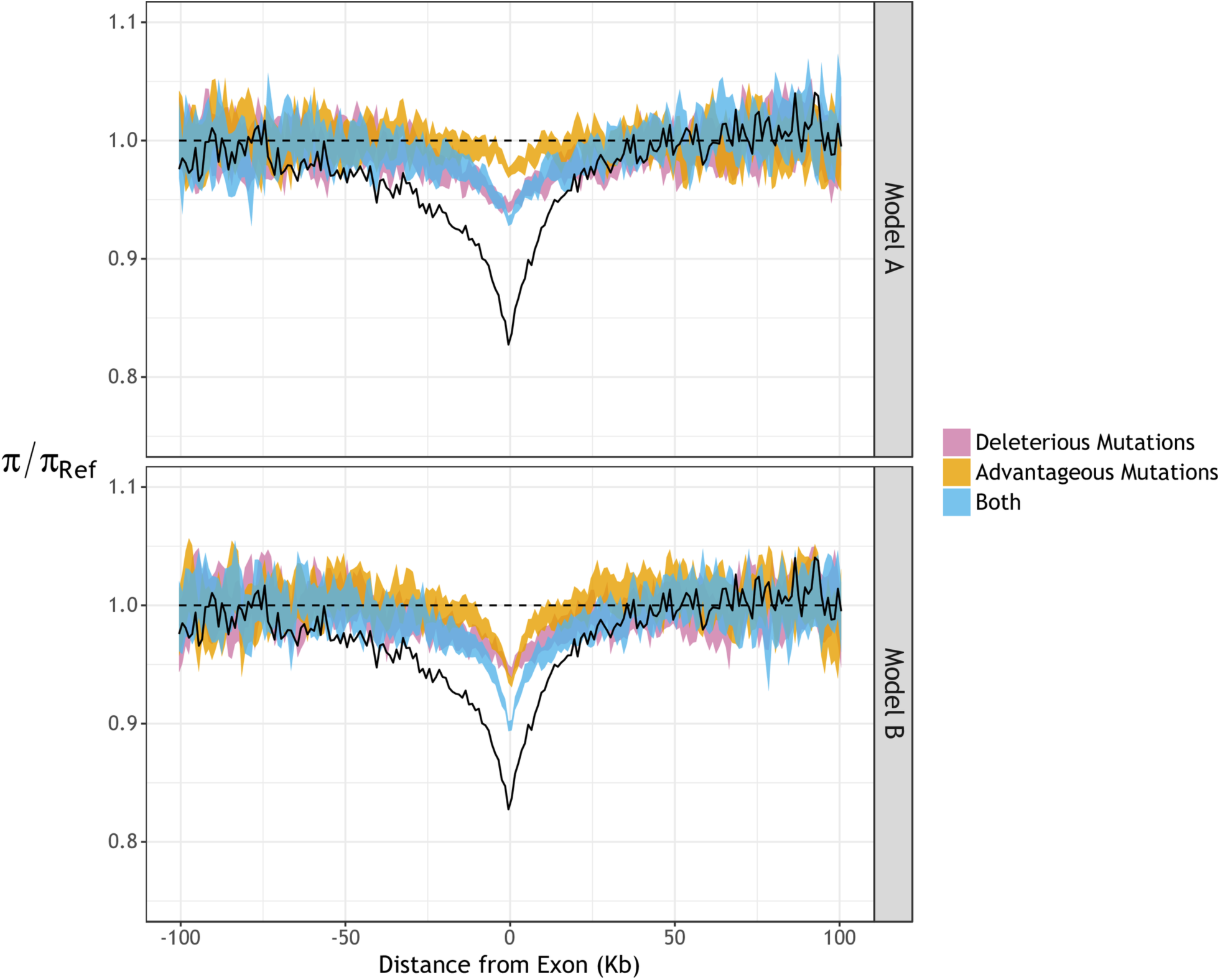
Estimates of scaled diversity (*π/π*_*Ref*_) around protein-coding exons (black lines) in *M. m. castaneus* compared to results from simulations (colored ribbons). The panels show diversity observed in simulated populations assuming DFE estimates obtained by analysis of the full uSFS (Model A) or when sites fixed for the derived allele do not influence selection parameters (Model B). Nucleotide diversity (*π*) is scaled by the mean diversity at distances more than 75 Kbp from exons (*π*_*Ref*_). Colored ribbons represent 95% confidence intervals obtained from 1,000 bootstrap samples.

In our simulations of exons and surrounding regions, recurrent SSWs produced troughs in diversity, but they were both narrower and shallower than those observed in the house mouse. However, the results are sensitive to the model used to estimate selection parameters (Figure 2; Table 3). Assuming the selection parameters estimated under Model A (i.e. analysing the full uSFS) we found that advantageous mutations produced a small dip in diversity around exons, which was shallower and narrower than the one generated by deleterious mutations alone (Figure 2; Table 3). In contrast, the advantageous mutation parameters estimated under Model B (i.e. where sites fixed for the derived allele do not influence selection parameters) resulted in a marked trough in diversity around exons in simulations (Figure 2; Table 3). In simulations that incorporated both advantageous and deleterious mutations, the troughs in diversity around exons were not as large as those observed in *M. m. castaneus* (Figure 2; Table 3). However, assuming Model B selection parameters resulted in a trough in diversity that was both deeper and wider than the one generated when assuming Model A parameters (Figure 2). The differences between Model A simulations and Model B simulations presumably arise because under Model B the frequency of advantageous nonsynonymous mutations was ∼3 times higher than under Model A (Table 2).

**Table 3.** The root mean square difference between values of *π* around functional elements predicted in simulations and *π* observed in *M. m. castaneus*. Confidence intervals were obtained from 1,000 bootstrap samples (*see Methods*).

|  |  | Exons |  | CNEs |  |
| --- | --- | --- | --- | --- | --- |
|  |  | Median | 95% range | Median | 95% range |
| **Model A** | *Deleterious Mutations* | 0.0327 | 0.0311 - 0.0343 | 0.0164 | 0.0151 - 0.0179 |
|  | *Advantageous Mutations* | 0.0422 | 0.0403 - 0.0442 | 0.0177 | 0.0161 - 0.0195 |
|  | *Both* | 0.0312 | 0.0297 - 0.0340 | 0.0100 | 0.0088 - 0.0113 |
| **Model B** | *Deleterious Mutations* | 0.0331 | 0.0314 - 0.0351 | 0.0157 | 0.0144 - 0.0171 |
|  | *Advantageous Mutations* | 0.0380 | 0.0355 - 0.0406 | 0.0162 | 0.0147 - 0.0179 |
|  | *Both* | 0.0274 | 0.0253 - 0.0294 | 0.0088 | 0.0078 - 0.0101 |

We also carried out simulations focussing on CNEs and found that the combined effects of BGS and recurrent SSWs, as generated by our estimates of selection parameters, can explain patterns of diversity observed in *M. m. castaneus* (Figure 3; Table 3). Selection parameters obtained under Models A and B produced similar results. The troughs in diversity around CNEs in simulations incorporating only advantageous mutations were similar to the ones generated by deleterious mutations alone (Figure 3; Table 3). Although both processes are required to explain the patterns observed in mice, our simulations suggest that BGS makes a bigger contribution to the overall reduction in neutral diversity than SSWs (Figure S3). The troughs in diversity around CNEs in our simulations were slightly shallower than those observed in the mouse genome (Figure 3), perhaps suggesting that we failed to detect infrequent, strongly selected advantageous mutations in CNEs or that we slightly underestimated the true frequency of advantageous mutations occurring in those elements.

**Figure 3.**
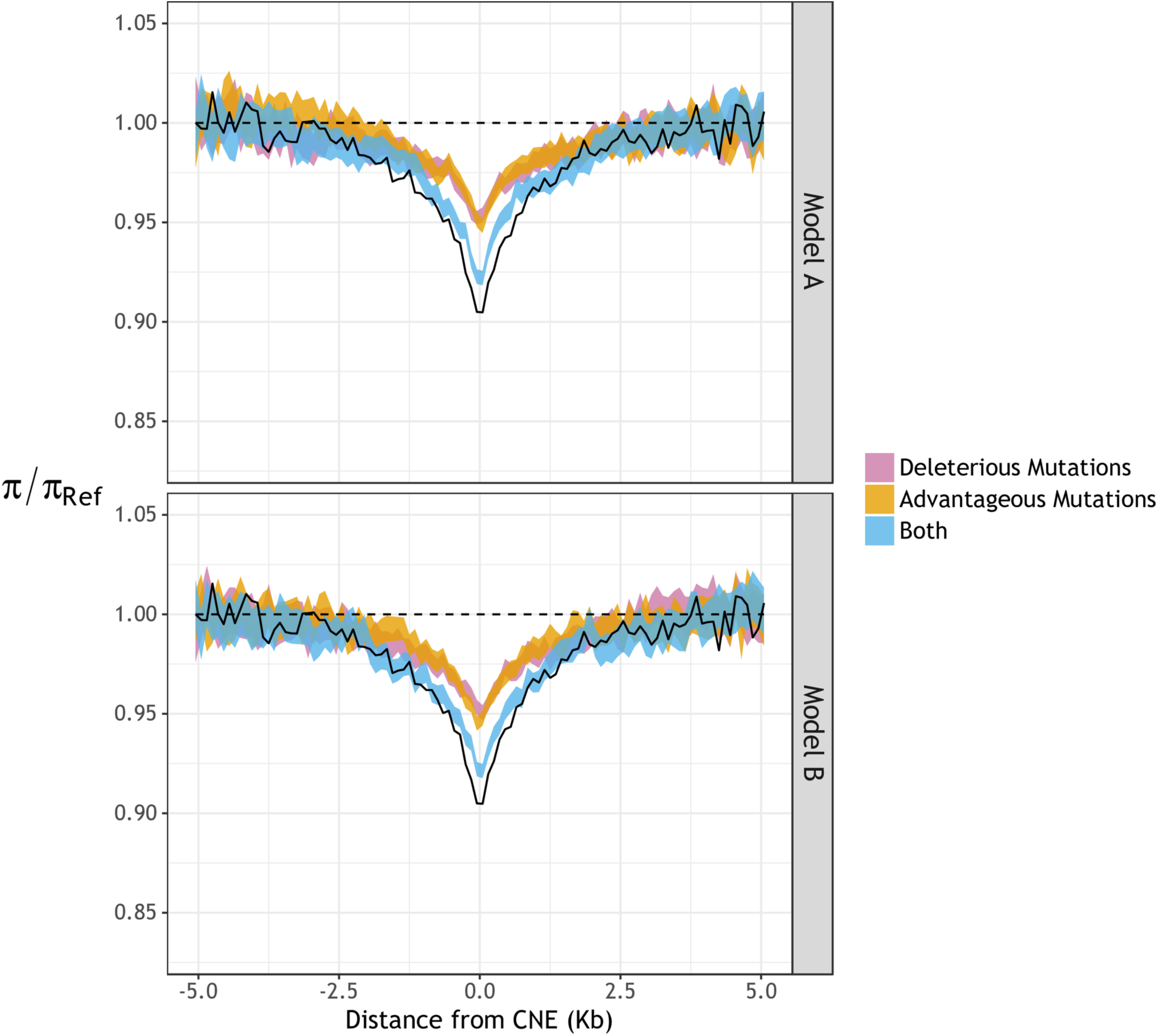
Estimates of scaled diversity (*π/π*_*Ref*_) around conserved non-coding elements (CNEs) in *M. m. castaneus* compared to results from simulations (colored ribbons). The panels show diversity observed in simulated populations assuming DFE estimates obtained by analysis of the full uSFS (Model A) or when sites fixed for the derived allele do not influence selection parameters (Model B). Nucleotide diversity (*π*) is scaled by the mean diversity at distances more than 4 Kbp from CNEs (*π*_*Ref*_). Colored ribbons represent 95% confidence intervals obtained from 1,000 bootstrap samples.

#### ii) The site frequency spectrum around functional elements

SSWs and BGS are known to affect the shape of the SFS for linked neutral sites [32, 33, 52]. SSWs and BGS generate troughs in diversity at linked sites (Figures 2–3), but nucleotide diversity on its own does not contain information about the shape of the SFS. Tajima’s *D* is a useful statistic for this purpose, because it is reduced when there is an excess of rare polymorphisms relative to the neutral expectation and increased when intermediate frequency variants are more common [53]. We therefore compared Tajima’s *D* in the regions surrounding functional elements in simulations with values observed in the real data. It is notable that average Tajima’s *D* is far lower in *M. m. castaneus* than in our simulations (Figure 4). This likely reflects a genome-wide process, such as population size change, that we have not modelled.

**Figure 4.**
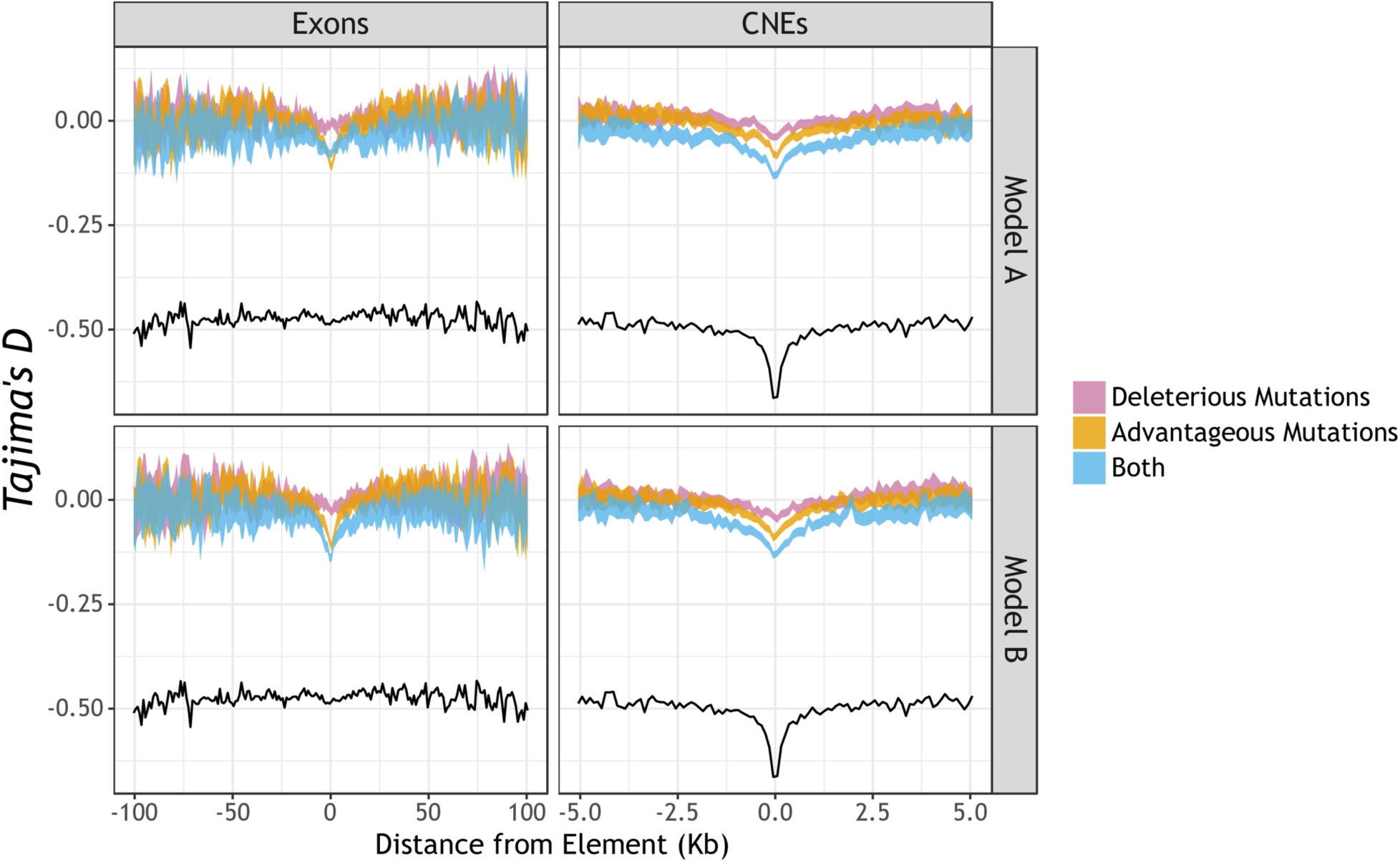
Tajima’s D around protein-coding exons and CNEs in *M. m. castaneus* compared to simulated data. The black lines show Tajima’s D computed from the *M. m. castaneus* genome sequence data around protein-coding exons or CNEs. The colored ribbons show the 95% bootstrap intervals from simulated data assuming the DFEs estimated under either Model A (i.e. analyzing the full uSFS) or Model B (i.e. fixed derived sites do not contribute to the likelihood for selection parameters).

If we assume selection parameters obtained under Model A, Tajima’s *D* around protein-coding exons is relatively invariant, and matches the pattern observed in the real data fairly well (Figure 4). However, under Model B, the simulations exhibit a substantial dip in Tajima’s *D*, which is not observed in the real data (Figure 4).

In the case of CNEs, we observed a trough in Tajima’s *D* in the real data (Figure 4), and simulations predict similar troughs under Models A and B (Figure 4). However, the trough in Tajima’s *D* may be caused by the presence of functionally constrained sequences in the immediate flanks of CNEs (*See Methods*), making a comparison between the simulations and the observed data problematic.

#### iii) Rates of substitution in functional elements

Incorporating information from sites fixed for the derived allele when estimating the DFE (as in Model A) or disregarding this information (as in Model B) had a striking effect on estimates of the frequency and effects of advantageous mutations (Table 2). In the case of 0-fold sites, for example, *p*_*a*_ was ∼3x higher under Model B than Model A (Table 2). We then investigated the extent by which such differences affect the divergence at selected sites under the two models. Nucleotide divergence at putatively neutral sites between the mouse and the rat is approximately 15%, so we simulated an expected neutral divergence of 7.5% for one lineage.

We compared the ratio of nucleotide divergence at selected sites to the divergence at neutral sites (*d*_*sel*_/*d*_*neut*_) between the simulated and observed data. In simulations that assumed the estimates of selection parameters obtained under Model A, *d*_*sel*_/*d*_*neut*_ values were similar to those observed in *M. m. castaneus* for all classes of selected sites (Table 4). Under Model B, however, the simulations predicted substantially more substitutions at nonsynonymous sites and UTRs than were seen in the real data (Table 4). This suggests that, under Model B, *p*_*a*_ for 0-fold sites and UTRs may be overestimated.

**Table 4.**
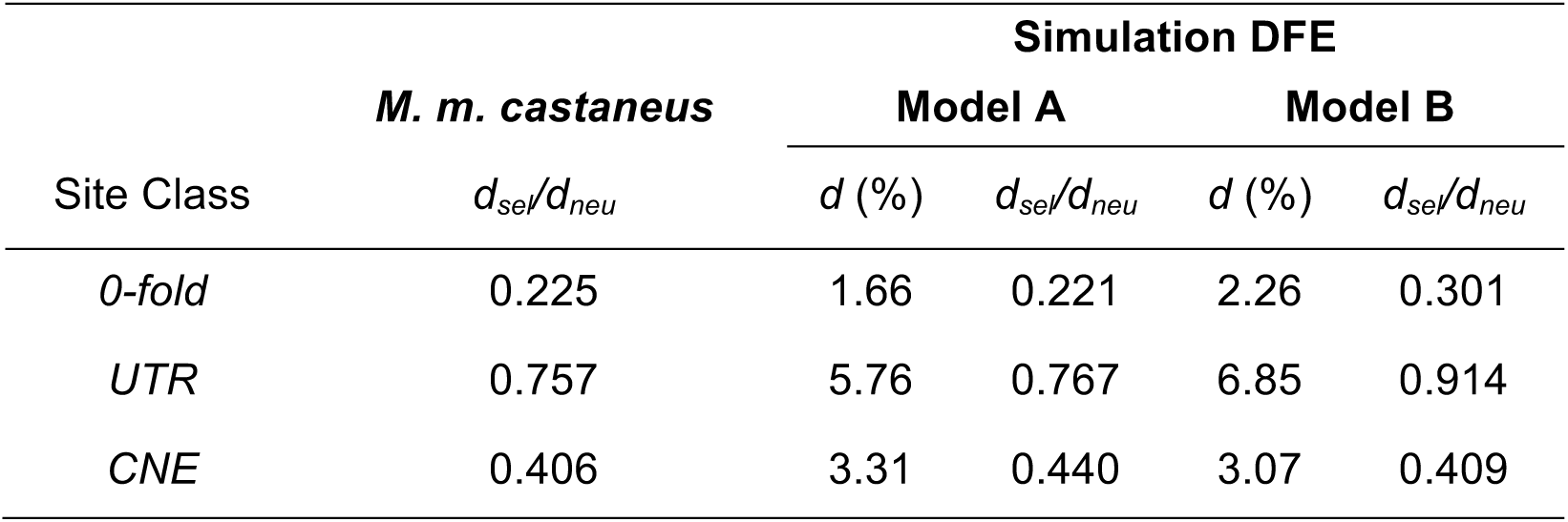
Comparison of the accumulation of nucleotide divergence in simulated populations between different functional site types. In the cases of 0-fold sites and UTRs, *d*_*neu*_ refers to 4-fold sites. For CNEs, *d*_*neu*_ refers to CNE flanking sites. In all simulations, *d*_*neu*_ was set to 7.5%.

#### iv) Re-estimating the DFE from simulated data

BGS and SSWs both perturb allele frequencies at linked neutral sites, and this can lead to the inference of spurious demographic histories [45–47]. By fitting a model incorporating three epochs of population size to the putatively neutral site data, we inferred that *M. m. castaneus* has experienced a population bottleneck followed by an expansion (Table S2). To investigate the possibility that the inferred demographic histories could be an artefact of selection at linked sites, we fitted demographic models to the uSFS obtained from simulated synonymous sites. Simulations assumed the selection parameters obtained under either Model A or B, and in each case, the 3-epoch model gave the best fit to the data. The estimated demographic parameters inferred were somewhat different between simulations assuming Model A or Model B selection parameters, but in each case a population bottleneck followed by an expansion was inferred (Table S5). This is an interesting observation, since our simulations assumed a constant population size, but selection at linked sites appears to distort the neutral site uSFS, and a demographic history is estimated as the one inferred from the real data (Table S5).

Our simulations also indicate that selection parameters are difficult to accurately infer using the uSFS alone. In the case of Model A simulations, the selection strength and frequency of deleterious mutations was accurately estimated, as was the combined frequency of all effectively neutral mutations (Table S5). However, in Model A simulations, DFE-alpha did not accurately estimate the strength and frequency of advantageous mutations. Estimates of selection parameters in Model B simulations were similar to the input parameters, but a notable exception was that the frequency of advantageous mutations (*p*_*a*_) was overestimated (Table S5). A possible explanation for this is that the demographic correction we applied to the uSFS for selected sites (see Supplementary Methods) may not fully capture the effects of selection at linked sites. SSWs increase the proportions of high frequency derived alleles [32], and it is possible that their contribution to the uSFS for selected sites was partially unaccounted for, creating the appearance of more frequent advantageous mutations in the uSFS.

#### v) Patterns of diversity around sites that have recently experienced a substitution

In general, it has been difficult to discriminate between BGS and SSWs, because their effects on genetic diversity and the site frequency spectrum are qualitatively similar. One method that has been suggested as a means of teasing the two processes apart takes advantage of the fact that hard SSWs should be centred on a nucleotide substitution, whereas this is not the case for BGS. Comparing the average genetic diversity in regions surrounding recent putatively selected and putatively neutral substitutions (e.g. 0-fold and 4-fold sites, respectively) may therefore reveal the action of SSWs [17, 19]. Halligan *et al.* [20] performed such an analysis in *M. m. castaneus* using the closely related *M. famulus* as an outgroup, and found that the profiles of neutral diversity around 0-fold and 4-fold substitutions were virtually identical. Similar findings have been reported in other species [19, 21]. One interpretation of these results is that hard SSWs are rare. To investigate this, we measured the average neutral diversity around nonsynonymous and synonymous substitutions in simulations for the case of frequent hard SSWs.

In our simulations, we measured diversity around substitutions occurring on a time-scale that is equivalent to the divergence time between *M. m. castaneus* and *M. famulus*. The average diversities around nonsynonymous and synonymous substitutions in the simulated data were very similar, regardless of whether simulations assumed the selection parameters estimated under Model A or Model B (Figure 5). However, the troughs in diversity around substitutions were deeper in the simulations assuming Model B (Figure 5), reflecting the higher frequency of advantageous mutations (Table 2). In the immediate vicinity of nonsynonymous substitutions, diversity was lower than the corresponding value for synonymous substitutions (Figure 5). However, the differences are slight, so it would be difficult to draw firm conclusions about the action of either SSWs or BGS. Taken together, these results suggest that analysing patterns of diversity around recent substitutions does not provide enough information that can convincingly discriminate between SSWs and BGS in *M. m. castaneus*, even when hard sweeps are fairly frequent. Further analysis is required to assess whether this is also the case for other organisms.

**Figure 5.**
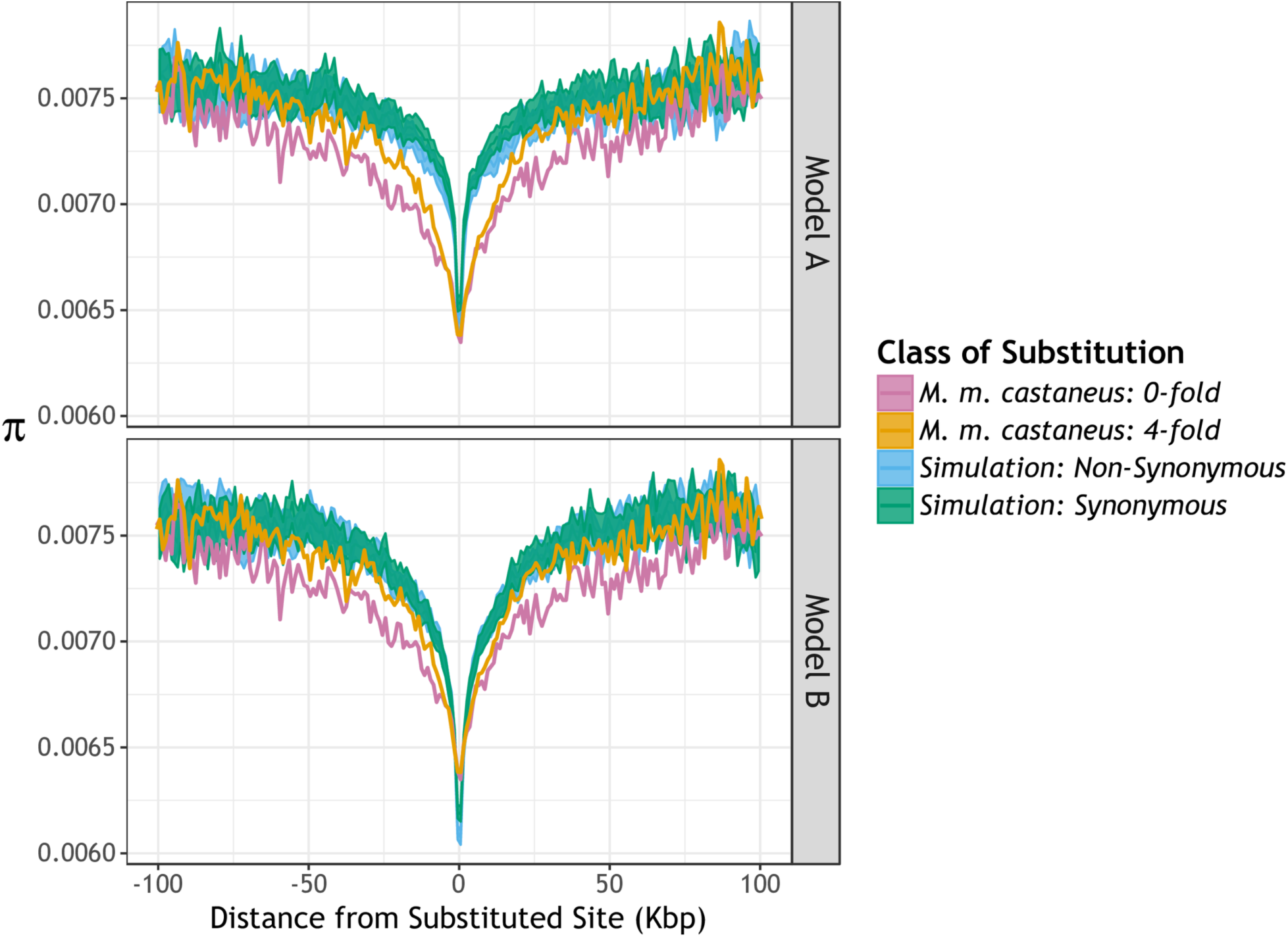
Nucleotide diversity (*π*) around substituted sites in *M. m. castaneus* compared to the same pattern obtained from simulation data. Nucleotide diversity in *M. m. castaneus* was scaled by divergence between mouse and rat to correct for variation in local mutation rates. The *M. m. castaneus* data are from [20].

## Discussion

There are a number of observations suggesting that natural selection is pervasive in the murid genome. First, there is a positive correlation between synonymous site diversity and the rate of recombination [44]. Secondly, there is reduced diversity on the X-chromosome compared to the autosomes, which cannot readily be explained by neutral or demographic processes [28]. Thirdly, there are troughs in genetic diversity surrounding functional elements, such as protein-coding exons and CNEs, which are consistent with the action of background selection (BGS) and/or SSWs [20]. In this paper, we analysed the genome sequences of 10 *M. m. castaneus* individuals sampled from the ancestral range of the species [20]. We estimated the DFEs for several classes of functional sites (0-fold nonsynonymous sites, UTRs and CNEs), and used these estimates to parameterise forward-in-time simulations. We investigated whether the simulations predict the observed troughs in diversity around functional elements along with the between-species divergence observed between mice and rats.

### Estimating selection parameters based on the uSFS

Relative to putatively neutral comparators, 0-fold sites, UTRs and CNEs all exhibit reduced nucleotide diversity, reduced nucleotide divergence and an excess of low frequency variants (Table 1; Figure 1), consistent with the action of natural selection [20, 27]. The estimates of the DFEs included substantial proportions of strongly deleterious mutations (Table 2). In addition, the best-fitting models also included a single class of advantageous mutations. Additional classes were not statistically supported, however. In reality, there is almost certainly a distribution of advantageous selection coefficients [54, 55]. A visual examination of the fitted and observed uSFSs, however, shows that the best-fitting DFEs fit the data very well (Figure S5), suggesting that there is limited information in the uSFS to estimate a range of positive selection coefficients.

When estimating the DFE for a particular class of sites, we analysed either the full uSFS including sites fixed for the derived allele (Model A) or we ignored sites fixed for the derived allele (i.e. Model B). Recently, Tataru *et al.* [26] used simulations to show that selection parameters can be accurately estimated from the uSFS, whilst ignoring between-species divergence, if *p*_*a*_ is sufficiently high. In our analysis of 0-fold sites and UTRs, Model B gave a significantly better fit and higher estimates of the frequency of advantageous mutations (*p*_*a*_) than Model A (Table 2). For CNEs, however, Models A and B did not significantly differ in fit, and the selection parameter estimates were very similar (Table 2). The goodness-of-fit and parameter estimates obtained under Models A and B may differ if the processes that generated between species-divergence are decoupled from the processes that produce within species diversity. There are several factors that could potentially cause this decoupling. 1) Past demographic processes may have distorted the uSFS in ways not captured by the corrections we applied; 2) there may be error in assigning alleles as ancestral or derived; 3) the nature of the DFE may have changed in the time since the accumulation of between-species divergence began; and 4) there could be rare, strongly advantageous mutations that contribute to divergence, but contribute negligibly to polymorphism. It is difficult to know which of these factors affected the outcome of our analyses. However, we found that Model B gave a better fit to the uSFS than Model A for 0-fold sites and UTRs, but not CNEs. In addition, we that found that the selection parameters obtained fail to explain the patterns of diversity around protein-coding exons, whereas they explain the patterns of diversity around CNEs, so we think the latter explanation is likely to have been important.

### Patterns of diversity and Tajima’s D around functional elements

We performed simulations incorporating our estimates of deleterious and advantageous mutation parameters to dissect the contribution of BGS and selective sweeps to patterns of diversity around functional elements. We found that BGS does not fully explain the troughs in diversity observed around either protein-coding exons or CNEs (Figures 2–3). These results are consistent with Halligan *et al.* [20].

Our simulations suggest that BGS and SSWs both produce genome-wide reductions in neutral diversity (Figures S3–4), but neither process on its own fully explains the troughs in diversity around protein-coding exons and CNEs, regardless of which model (A or B) is used to estimate selection parameters (Figures 2–3). Around protein-coding exons, the combined effects of advantageous and deleterious mutations generated a shallower trough in diversity than the one observed (Figure 2). A possible explanation for this is that rare, strongly selected advantageous mutations are undetectable by analyses based on the uSFS (discussed below). In contrast, the combined effects of BGS and SSWs predicted troughs in diversity surrounding CNEs that closely match those observed (Figure 3).

There is an overall excess of rare variants in *M. m. castaneus* relative to neutral expectation, as indicated by a strongly negative Tajima’s *D* at putatively neutral sites (Table 1) and in the regions surrounding exons and CNEs (Figure 4). Our simulations incorporating both advantageous and deleterious mutations also exhibited negative Tajima’s *D*, but not nearly so negative as in the real data (Figure 4). This difference between the observed data and the simulations indicates that there may be processes generating an excess of rare variants, such as a recent population expansion, which were not incorporated in the simulations.

### Rates of nucleotide substitutions in simulations

Our simulations suggest that the frequency of advantageous mutations (*p*_*a*_) estimated for 0-fold sites and UTRs under Model B may be unrealistically high. This is because several aspects of the results were incompatible with the observed data. Firstly, we found that the substitution rates for simulated nonsynonymous and UTR sites were higher than those observed between mouse and rat (Table 4). Secondly, we observed a pronounced dip in Tajima’s *D* around simulated exons, which is not present in the real data (Figure 4), suggesting that under Model B, either the strength or frequency of positive selection at 0-fold sites is overestimated.

### Do our results provide evidence for strongly selected advantageous mutations?

Estimation of the rate and frequency of advantageous mutations based on the uSFS relies on the presence of advantageous variants segregating within the population [23, 25, 26]. The frequency of advantageous mutations may impose a limit on the parameters of positive selection that can be accurately estimated. Indeed, Tataru *et al*. [26] recently showed that *p*_*a*_ may be overestimated when analysing the uSFS, if the true value of *p*_*a*_ is low.

Advantageous mutations with large effects have shorter sojourn times than those with milder effects [56, 57]. If strongly selected advantageous mutations are infrequent, it is therefore unlikely that they would be observed to be segregating. This could explain why the estimated selection parameters fail to predict the deep troughs in diversity around exons that we observe in the real data (Figure 2). Furthermore, the fact that Model B gave a better fit than Model A for 0-fold sites and UTRs suggests that polymorphism and divergence have become decoupled for those sites. This is also consistent with the presence of infrequent, strongly selected mutations that become fixed rapidly and are thus not commonly observed as polymorphisms.

Relevant to this point, an interesting comparison can be made between two recent studies to estimate the frequency and strength of positive selection using the same *D. melanogaster* dataset. The first, by Keightley *et al.* [31], utilised the uSFS analysis methods of Schneider *et al.* [25] (i.e. Model A in the present study), and estimated the frequency of advantageous mutations (*p*_*a*_) = 4.5 × 10^−3^ and the scaled strength of selection (*N*_*e*_*s*_*a*_) = 11.5 for 0-fold nonsynonymous sites. The second study, by Campos *et al*. [43], estimated *p*_*a*_ *=* 2.2 × 10^−4^ and *N*_*e*_*s*_*a*_ = 241, based on the correlation between synonymous site diversity and nonsynonymous site divergence. Although the individual parameter estimates differ substantially, the compound parameter *N*_*e*_*s*_*a*_*p*_*a*_ (which approximates the rate of SSWs) was similar between the studies (0.055 and 0.052 for Campos *et al*. [43] and Keightley *et al.* [31] respectively). It is expected that synonymous site diversity is reduced by SSWs, so the method used by Campos *et al*. [43] may be sensitive to the presence of strongly selected mutations, whereas the Keightley *et al.* [31] approach may have been more sensitive to weakly selected mutations. It seems plausible then, that the two studies capture different aspects of the DFE for advantageous mutations (a similar argument was made by Sella *et al.* [58]). Supporting this view, Elyashiv *et al.* [5] recently estimated the DFE in *D. melanogaster*, incorporating both strongly and weakly selected advantageous mutations, by fitting a model incorporating BGS and SSWs to genome-wide variation in genetic diversity. They inferred that weakly selected mutations are far more frequent than strongly selected ones. In the present study, we used similar methods as Keightley *et al.* [31] to estimate the frequency and strength of advantageous mutations, so the estimated parameters of positive selection may represent only weakly selected mutations. Indeed, patterns of diversity at microsatellite loci suggest that there are strongly selected, infrequent sweeps in multiple European *M. musculus* populations [59], so infrequent strong sweeps may be a general feature of mouse evolution.

The patterns of diversity and Tajima’s *D* around CNEs and the nucleotide divergence within CNEs in our simulated populations were similar to those observed in the *M. m. castaneus* data, regardless of which estimate of the DFE we used (i.e. Model A or B) (Figure 3–4; Table 3). This suggests that the four classes of mutational effects inferred provide a reasonable approximation for the full distribution of fitness effects for CNEs.

Understanding the contributions of regulatory and protein change to phenotypic evolution has been an enduring goal in evolutionary biology [60–62]. If selection is strong relative to drift (i.e. *N*_*e*_*s*_*a*_ > 1) then the rate of change of fitness due to advantageous mutations is expected to be proportional to the square of the selection coefficient [63]. In this study, we inferred that the strength of selection acting on new advantageous mutations in CNEs and 0-fold sites are roughly equivalent, but that advantageous mutations occur more frequently in CNEs (Table 2). Given that there are more CNE nucleotides in the genome than there are 0-fold sites (Table 1), this could imply that adaptation at regulatory sites causes the greatest fitness change in mice. However, we have argued that protein-coding genes may be subject to strongly selected advantageous mutations, which were undetectable by analysis of the uSFS. If this were the case, adaptation in protein-coding genes could make a larger contribution to fitness change than regulatory sites.

### Limitations of the study

There is a growing body of evidence suggesting that hard sweeps may not be the primary mode of adaptation in both *D. melanogaster* and humans. Firstly, soft sweeps, where multiple haplotypes reach fixation due to the presence of multiple *de novo* mutations or selection acted on standing variation, may be common. Garud *et al.* [64] developed a suite of haplotype-based statistics that can discriminate between soft and hard SSWs. The application of these statistics to North American and Zambian populations of *D. melanogaster* suggested that soft sweeps are the dominant mode of adaptation in that species, at least in recent evolutionary time [64, 65]. Furthermore, Schrider and Kern [66] recently reported that signatures of soft sweeps are more frequent than those of hard sweeps in humans. However, their method did not explicitly include the effects of partial sweeps and/or BGS. Under a model of stabilising selection acting on a polygenic trait, if the environment changes, adaptation to a new optimum may cause small shifts in allele frequency at numerous loci without necessarily resulting in fixations [67, 68]. Genome-wide association study hits in humans exhibit evidence that such partial SSWs may be common [69]. These results all suggest that the landscape of adaptation may be more complex than the model of directional selection acting on a *de novo* mutation assumed in this study. For example, our simulations did not incorporate changing environments or stabilising selection, so we were unable to model adaptive scenarios other than hard sweeps.

Further work should aim to understand the probabilities of the different types of sweeps. Different functional elements have different DFEs for harmful mutations. In particular, regulatory elements seem to experience more mildly selected deleterious mutations than coding sequences [18, 20] (Table 2). It has been argued that such differences in constraint between coding and non-coding elements may be due to a lower pleiotropic burden on regulatory sequences [61]. Differences in the DFE among different genomic elements is expected to affect genetic diversity within these elements. This, in turn, may affect the modes of sweeps that occur, since the relative probabilities of a hard or soft sweep depend on the level of standing genetic variation (reviewed in [70]).

In our simulations, we treated *N*_*e*_ as constant through time, but this is likely to be an oversimplification. We analysed two different classes of putatively neutral sites, and inferred there has been a population size bottleneck followed by an expansion (Table S2). In our simulations, however, we showed that the inferred demographic history may largely be an artefact of selection at linked sites (Table S5). There is a strongly negative Tajima’s *D* in genomic regions far from functional elements, which is not explained by selection (or at least the selection parameters we inferred) (Figure 4). This reduction is presumably caused by a demographic history or strong selection that was not included in our simulations. Less biased estimates of the demographic history of *M. m. castaneus* may be obtained from regions of the genome experiencing high recombination rates, located far from functional elements. Finally, mouse populations may rapidly oscillate in size (e.g. seasonally [71]). If this were the case, so would the effective selection strength of new mutations (and thus the probabilities of SSWs) [72].

In house mice, crossing over events predominantly occur in narrow windows of the genome termed recombination hotspots [73]. The locations of recombination hotspots have evolved very rapidly between and within *M. musculus* sub-species [74]. Assuming a single suite of recombination hotspots in simulations may produce misleading results if hotspot locations evolve faster than the rate of neutral coalescence. Recombination hotspots are an important feature of the recombination landscape in mice and thus potential influence the patterns of diversity around functional elements, but the appropriate way to model them is unclear.

## Conclusions

Using simulations, we have shown that estimates of the DFE obtained by analysis of the uSFS can explain the patterns of diversity around CNEs, but not around protein-coding exons. We also argue that mutations with moderately advantageous effects frequently occur at 0-fold and UTR sites, but that undetectable, strongly advantageous mutations may occur in both these classes of sites. Estimates of the strength and rate of advantageous mutations could be obtained by directly fitting a sweep model to the troughs in diversity around functional elements. We have shown that BGS makes a substantial contribution to these troughs, and using models that incorporate both BGS and sweeps [5, 43, 75] might allow us to make more robust estimates of selection parameters.

## Acknowledgements

We owe thanks to Brian Charlesworth for useful comments on the manuscript and to Deborah Charlesworth, Dan Halligan and the evolutionary genetics lab group at the University of Edinburgh for helpful discussions. Tom Booker is supported by a BBSRC EASTBIO Studentship. This project has received funding from the European Research Council (ERC) under the European Union’s Horizon 2020 research and innovation program (grant agreement No. 694212).

---

**Table S1.** Comparison of the fit of demographic models based on the analysis of 4-fold sites and CNE-flanks in *M. m. castaneus*.

| | Epochs | $\Delta \ln L$ | $\chi^2$ | # Estimated Parameters |
| --- | --- | --- | --- | --- |
| *4-fold* | 1 | 1,620 | 22,500 | 2 |
|  | 2 | 159 | 2,930 | 4 |
|  | 3 | 0.0 | 553 | 6 |
| *CNE-flank* | 1 | 19,100 | 53,500 | 2 |
|  | 2 | 1,350 | 5,070 | 4 |
|  | 3 | 0.0 | 975 | 6 |

**Table S2.** Parameters of the best-fitting demographic model estimated from the analysis of 4-fold and CNE-flanking sites.

|  | **4-fold** | **CNE-flank** |
| --- | --- | --- |
| $N_2/N_1$ | 0.40 | 0.07 |
| $t_2/N_1$ | 0.44 | 0.17 |
| $N_3/N_1$ | 0.40 | 1.00 |
| $t_3/N_1$ | 1.10 | 0.63 |

**Table S3.** Likelihood differences between models of the deleterious DFE (dDFE) fitted with or without a single class of adaptive mutations.

| Site Type | dDFE Model | $\Delta \ln L$ | |
| --- | --- | --- | --- |
|  |  | dDFE | dDFE + Adaptive Mutations |
| *0-fold* | 1-Class | 49,300 | 4.18 |
|  | 2-Class | 129 | **0.00** |
|  | 3-Class | 129 | 0.00 |
|  | Gamma | 247 | 4.18 |
| *CNE* | 1-Class | 51,000 | 245 |
|  | 2-Class | 1,660 | 3.41 |
|  | 3-Class | 1,480 | **0.00** |
|  | Gamma | 2,310 | 19.3 |
| *UTR* | 1-Class | 6,170 | 32.7 |
|  | 2-Class | 335 | **0.00** |
|  | 3-Class | 335 | 0.00 |
|  | Gamma | 970 | 13.5 |

**Table S4.** Parameter estimates for the scaled effect and frequency of advantageous mutations in three classes sites in *Mus musculus castaneus* when models incorporated either one class of advantageous mutations, or two.

| Number of<br>Advantageous<br>Mutation<br>Classes | 0-fold |  | UTR |  | CNE |  |
| --- | --- | --- | --- | --- | --- | --- |
|  | 1 | 2 | 1 | 2 | 1 | 2 |
| **Model A: Full uSFS** |  |  |  |  |  |  |
| $N_e s_a (1)$ | 7.27 | 7.27 | 5.32 | 5.32 | 9.17 | 9.17 |
| $p_a (1)$ | 0.003 | 0.003 | 0.0133 | 0.0133 | 0.0098 | 0.0098 |
| $N_e s_a (2)$ | - | 0.000 | - | 0.000 | - | 0.000 |
| $p_a (2)$ | - | 0.000 | - | 0.000 | - | 0.000 |
| $\Delta \ln L$ | - | 0.000 | - | 0.005 | - | 0.000 |
| **Model B: Sites fixed for the derived allele do not contribute to parameter estimates** |  |  |  |  |  |  |
| $N_e s_a (1)$ | 8.30 | 8.30 | 6.96 | 6.96 | 8.60 | 8.60 |
| $p_a (1)$ | 0.010 | 0.010 | 0.0294 | 0.0294 | 0.008 | 0.008 |
| $N_e s_a (2)$ | - | 0.0925 | - | 33.6 | - | 0.240 |
| $p_a (2)$ | - | 0.000 | - | 0.000 | - | 0.000 |
| $\Delta \ln L$ | - | 0.000 | - | 0.000 | - | 0.002 |

**Table S5.** Parameters of the selection model (2-Class dDFE + adaptation) when estimated from simulated data.

|  |  | Model A |  | Model B |  |
| --- | --- | --- | --- | --- | --- |
|  |  | Simulated Value | Estimated | Simulated Value | Estimated |
| | $N_e s (0)$ | -0.045 | -0.70 | -0.171 | -0.40 |
| | $p (0)$ | 0.191 | 0.145 | 0.184 | 0.181 |
| | $N_e s (1)$ | -104 | -92.3 | -100 | -77.3 |
| | $p (1)$ | 0.806 | 0.784 | 0.806 | 0.799 |
| | $N_e s (a)$ | 7.27 | 0.950 | 8.30 | 4.91 |
| | $p (a)$ | 0.00300 | 0.0710 | 0.0100 | 0.0200 |
| *Estimated in M. m. castaneus* |  |  |  |  |  |
| $N2/N1$ | 0.40 | - | 0.20 | - | 0.12 |
| $N3/N1$ | 1.4 | - | 1.0 | - | 0.9 |
| $t2/N2$ | 0.31 | - | 0.46 | - | 1.2 |
| $t3/N3$ | 0.79 | - | 1.4 | - | 1.3 |

**Figure S1.**
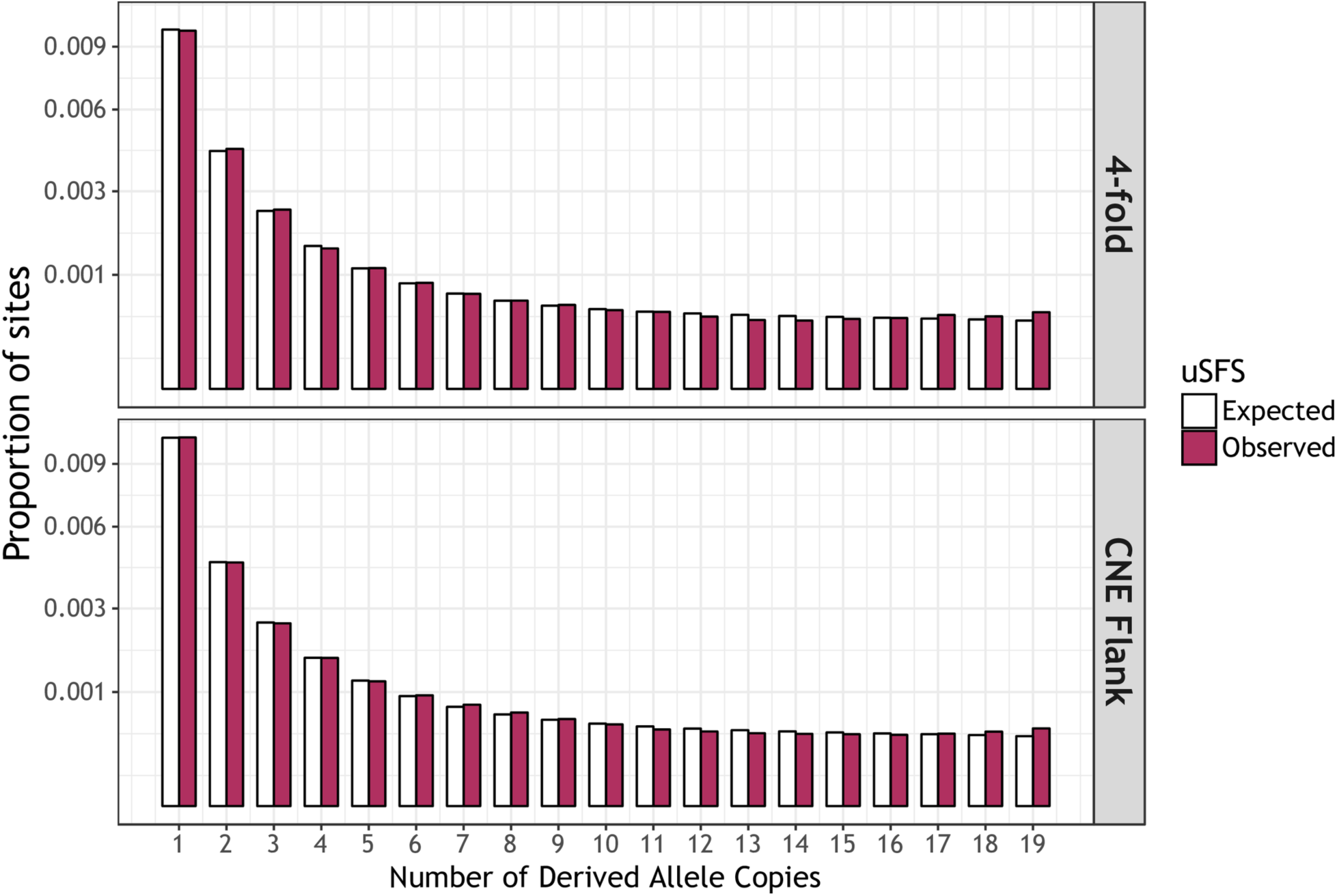
A comparison of the uSFS expected and observed under the best-fitting demographic models for two classes of putatively neutral sites, 4-fold degenerate synonymous sites and CNE-flanking sequences.

**Figure S2.**
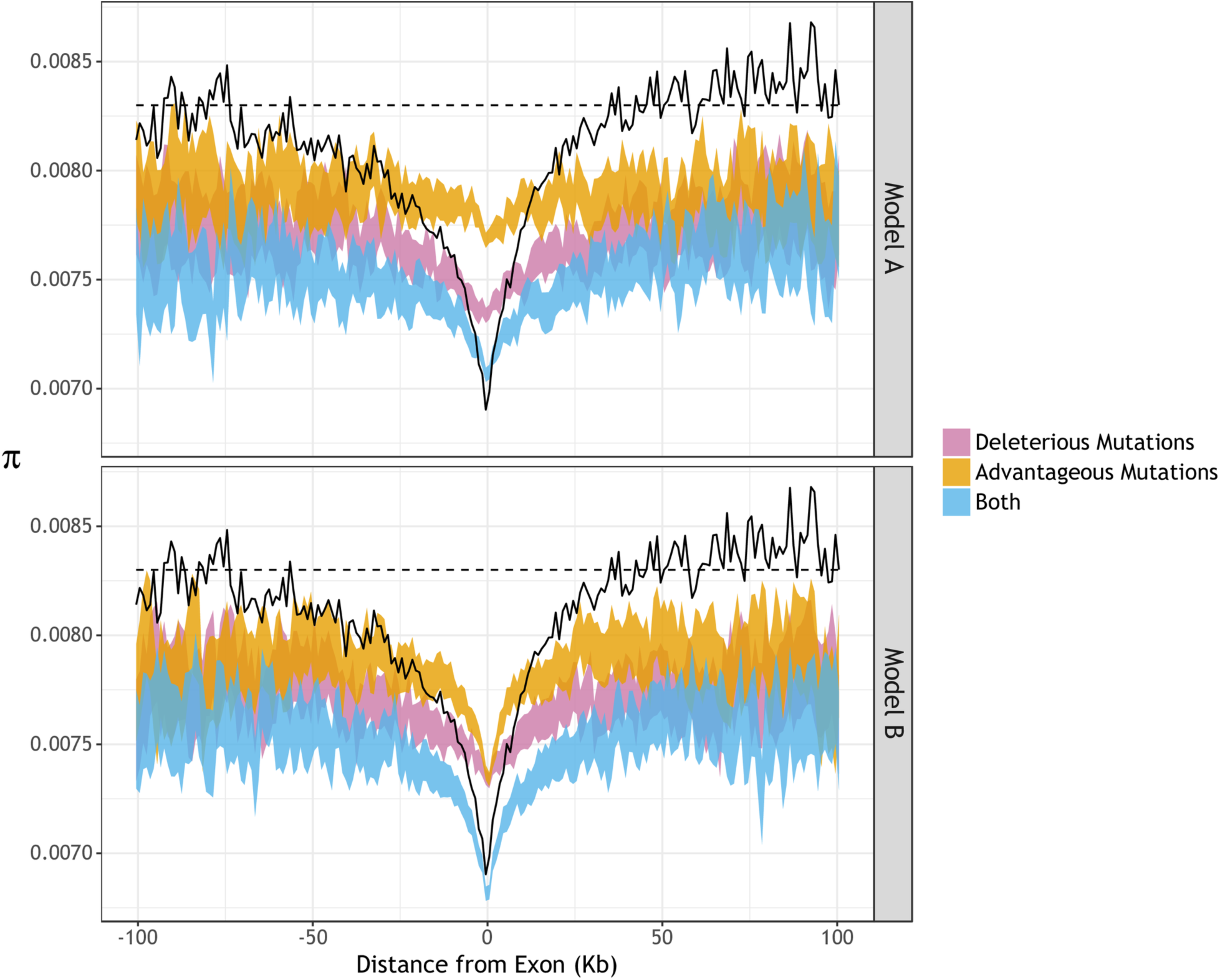
Estimates of unscaled *π* around protein-coding exons in *M. m. castaneus* (black line) compared to the values observed in simulated populations (coloured ribbons).

**Figure S3.**
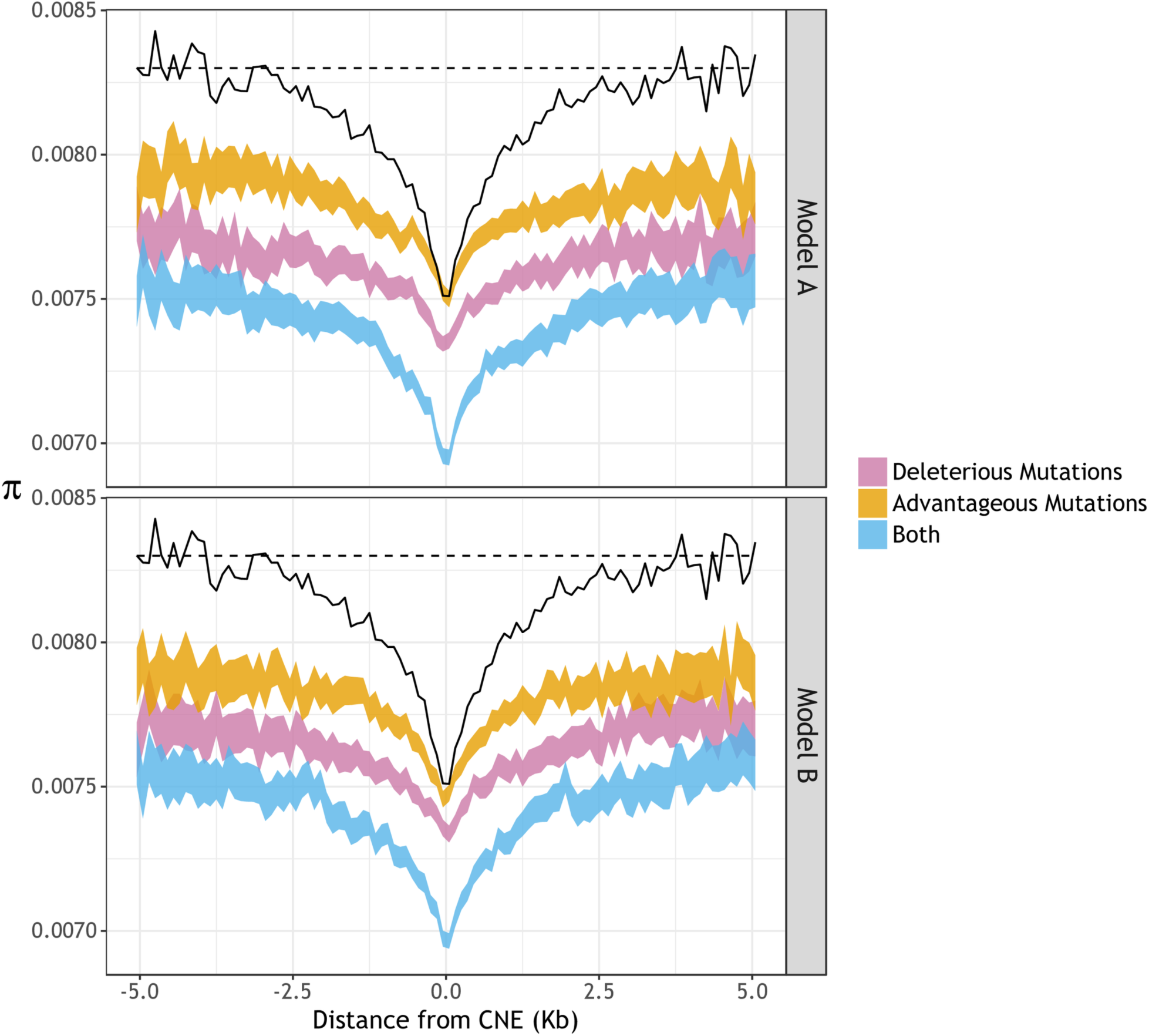
Estimates of unscaled *π* around conserved non-coding elements in *M. m. castaneus* (black line) compared to the values observed in simulated populations (coloured ribbons).

**Figure S4.**
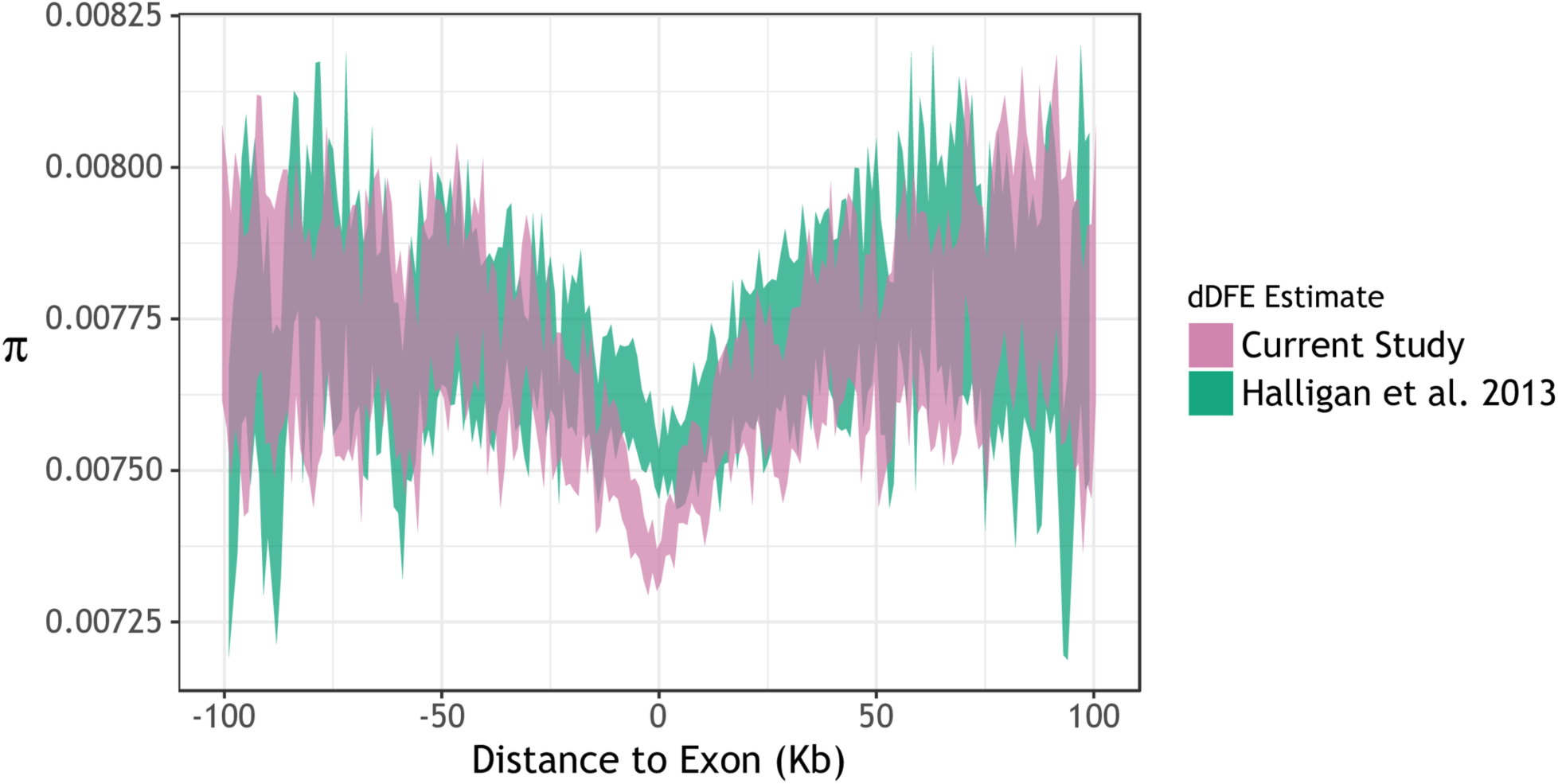
Comparison of nucleotide diversity (*π*) around protein-coding exons in simulated populations under either the discrete-class dDFE estimated in the current study or the gamma dDFE estimated by Halligan *et al.* [20].

**Figure S5.**
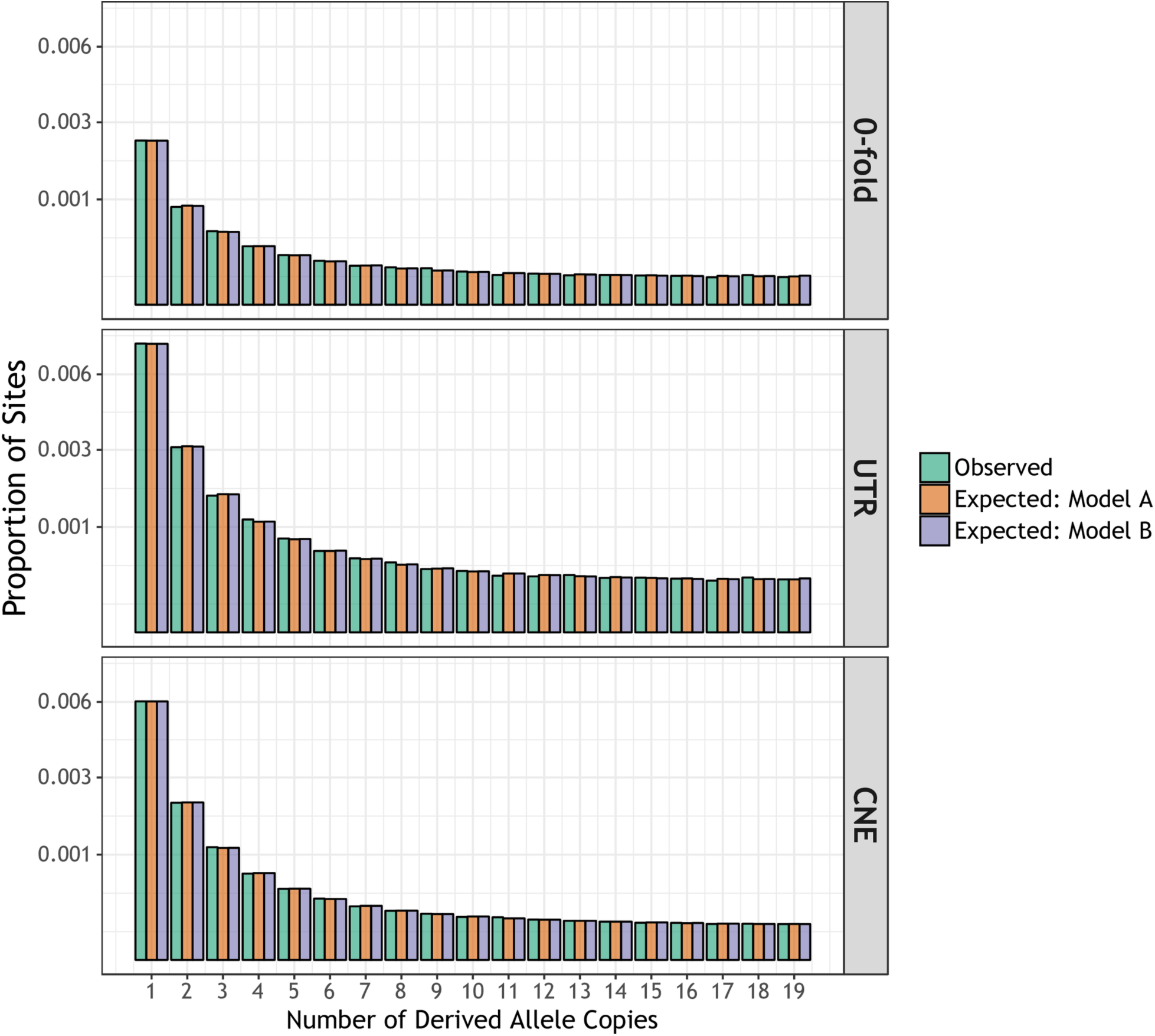
A comparison of the uSFS expected and observed under the best-fitting selection models for three classes of functional sites.

